# IFITM3 regulates virus-induced inflammatory cytokine production by titrating Nogo-B orchestration of TLR responses

**DOI:** 10.1101/2021.07.23.453513

**Authors:** M. Clement, J.L. Forbester, M. Marsden, P. Sabberwal, D. Wellington, S. Dimonte, S. Clare, K. Harcourt, Z. Yin, L. Nobre, R Antrobus, B. Jin, M. Chen, S. Makvandi-Nejad, J.A Lindborg, S.M. Strittmatter, M.P. Weekes, R.J. Stanton, T. Dong, I.R. Humphreys

## Abstract

Interferon induced transmembrane protein 3 (IFITM3) is an important viral restriction factor in viral pathogenesis that also exhibits poorly understood immune regulatory functions. Here, using human and mouse models, we demonstrate that IFITM3 regulates MyD88-dependent TLR-mediated cytokine production following dendritic cell exposure to cytomegalovirus (CMV), and this process limits viral pathogenesis *in vivo*. IFITM3 also restricted pro-inflammatory (IL-6) cytokine production in response to influenza. IFITM3 bound to and promoted ubiquitination and proteasomal degradation of the reticulon 4 isoform Nogo-B. We reveal that Nogo-B mediates TLR-dependent pro-inflammatory cytokine production and promotes viral pathogenesis *in vivo*, and this process involved alteration of TLR dynamics. The anti-inflammatory function of IFITM3 was intrinsically linked to its ability to regulate Nogo-B. Thus, we uncover Nogo-B as an unappreciated driver of viral pathogenesis and highlight a novel immune regulatory pathway where IFITM3 fine-tunes TLR responsiveness of myeloid cells to viral stimulation.

## Introduction

Interferon (IFN) induced transmembrane protein-3 (IFITM3) plays a major role in antiviral cellular defence, directly limiting cellular entry of a number of pathogenic viruses, including influenza A virus (IAV), HIV, vesicular stomatitis virus (VSV) and SARS-CoV (Everitt et al., 2012; Huang et al., 2011; Lu et al., 2011; Weidner et al., 2010). Some studies have demonstrated that IFITM3, along with other members of the IFITM family of proteins, may decrease cell membrane fluidity, possibly affecting viral fusion (Li et al., 2013; Lin et al., 2013). In one study, alteration in membrane fluidity was attributed to upregulation of cellular cholesterol (Amini-Bavil-Olyaee et al., 2013). However, subsequent studies have failed to demonstrate a mechanistic link between cholesterol levels and IFITM3 activity (Desai et al., 2014; Lin et al., 2013; Wrensch et al., 2014). IFITM3 has also been shown to act downstream of viral attachment, endocytosis and subsequent viral hemifusion, directly restricting full viral fusion and trapping virus particles in endosomes (Suddala et al., 2019). IFITM3 may also prevent virus entry by altering rates of virus-endosome fusion and/or accelerating the trafficking of endosomal cargo to lysosomes for destruction (Spence et al., 2019). Furthermore, IFITM3 and other IFITM proteins can directly interact with viral proteins such as HIV-1 Env and inhibit their processing, restricting virus fusion with target host cell membranes (Yu et al., 2015). Thus, IFITM3 can exert antiviral effects by influencing a broad range of cellular mechanisms.

Genetic polymorphisms within the IFITM3 locus have been linked to increased pathogenesis of viral infections such as IAV and SARS-CoV-2 (Wellington et al., 2019; Zhang et al., 2020). *IFITM3* SNP rs34481144, which is located in the *IFITM3* promoter region, is associated with lower *IFITM3* mRNA levels and has a strong association with disease severity in influenza cohorts. The substitution of the majority G allele with the minority C allele reduces IFITM3 expression levels, with reduced number of antiviral CD8^+^ T cells in IAV-infected lung tissue (Allen et al., 2017). The *CC* genotype of the rs12252 SNP, located in exon 1 of *IFITM3*, has been associated with increased severity of IAV infections in multiple studies (Everitt et al., 2012; Makvandi-Nejad et al., 2018).

The implication from human genetic studies is that increased viral pathogenesis in individuals with reduced IFITM3 abundance and/or function reflects only reduced control of viral entry into cells. Interestingly, however, severe IAV disease in individuals with the *CC* genotype of the rs12252 SNP is associated with high CCL2 levels that drive pathogenic monocyte responses and hypercytokinemia characterized by elevated levels of cytokines including IL-6 (Everitt et al., 2012; Zhang et al., 2013). This hypercytokinemia is associated with fatal H7N9 infection in individuals with the rs12252 *CC* genotype (Wang et al., 2014). Similarly, in mouse models of viral infection, Ifitm3 deficiency alters cytokine and chemokine profiles and leukocyte influx to sites of infection (Everitt et al., 2013; Gorman et al., 2016; Poddar et al., 2016). Importantly, in the murine cytomegalovirus (MCMV) model of infection, Ifitm3 restricts infection-induced IL-6 mediated viral disease, lymphopenia and loss of NK and T cells without directly impacting CMV replication (Stacey et al., 2017). In Sendai virus infection, IFITM3 inhibits IFN-β induction by promoting degradation of IRF3 within IFITM3-associated autophagosomes (Jiang et al., 2018). Moreover, mice lacking Ifitm1-3 produce elevated pro-inflammatory cytokines in response to chronic Poly I:C stimulation (Patoine et al., 2017). Thus, the biology of how IFITM3 limits viral pathogenesis may be complex and requires a better understanding.

Herein, we investigated how IFITM3 regulates virus-induced inflammatory responses using a combination of mice and human models of cytomegalovirus (CMV) infection, where viral entry is not susceptible to IFITM3-mediated entry restriction (Stacey et al., 2017; Xie et al., 2015), thus enabling dissection of the immune regulatory properties of IFITM3 independently of any direct antiviral restriction properties. We reveal that IFITM3 limits MyD88-dependent viral pathogenesis and that its anti-inflammatory properties are intrinsically linked with its ability to regulate the reticulon 4 isoform, Nogo-B. IFITM3 promoted Nogo-B turnover via ubiquitination and proteasomal degradation. We demonstrate in mice, and in human cellular models, that Nogo-B promotes TLR- and virus-induced inflammatory cytokine production and, in mice, pathogenesis. Finally, using human TLR2 as a model, we show that IFITM3-Nogo-B interactions alter TLR dynamics post-viral exposure, highlighting a potential mechanism for IFITM3-Nogo-B regulation of the inflammatory response during viral infection.

## Results

### Ifitm3 regulates MyD88-dependent TLR-mediated cytokine production following MCMV infection

We have previously demonstrated that Ifitm3 restricts IL-6 mediated MCMV pathogenesis *in vivo*, which associated with the limitation of cytokine production by DCs, but was accompanied by no defect in the direct control by Ifitm3 of virus replication (Stacey et al., 2017). To first provide evidence for a DC-intrinsic role for Ifitm3 in limiting viral disease, we irradiated wt mice and transferred bone marrow from zDC-DTR mice that enable conventional DC (cDC), depletion (Meredith et al., 2012) at a 50:50 ratio with *Ifitm3^-/-^* bone marrow. Mice were either treated (or not) with diphtheria toxin (DT) to generate mice that lacked Ifitm3 expression by DCs. Mice lacking Ifitm3^+^ cDCs (DT-depleted mice) demonstrated increased weight loss (Fig. 1A) as compared to mice with intact Ifitm3^+^ DCs. Exacerbated weight loss in mice lacking Ifitm3 expression by DCs was accompanied by elevated IL-6 production (Fig. 1B) without any impact on virus load (Fig. 1C), thus suggesting the importance of Ifitm3 expression by DCs in MCMV pathogenesis.

**Figure 1.**
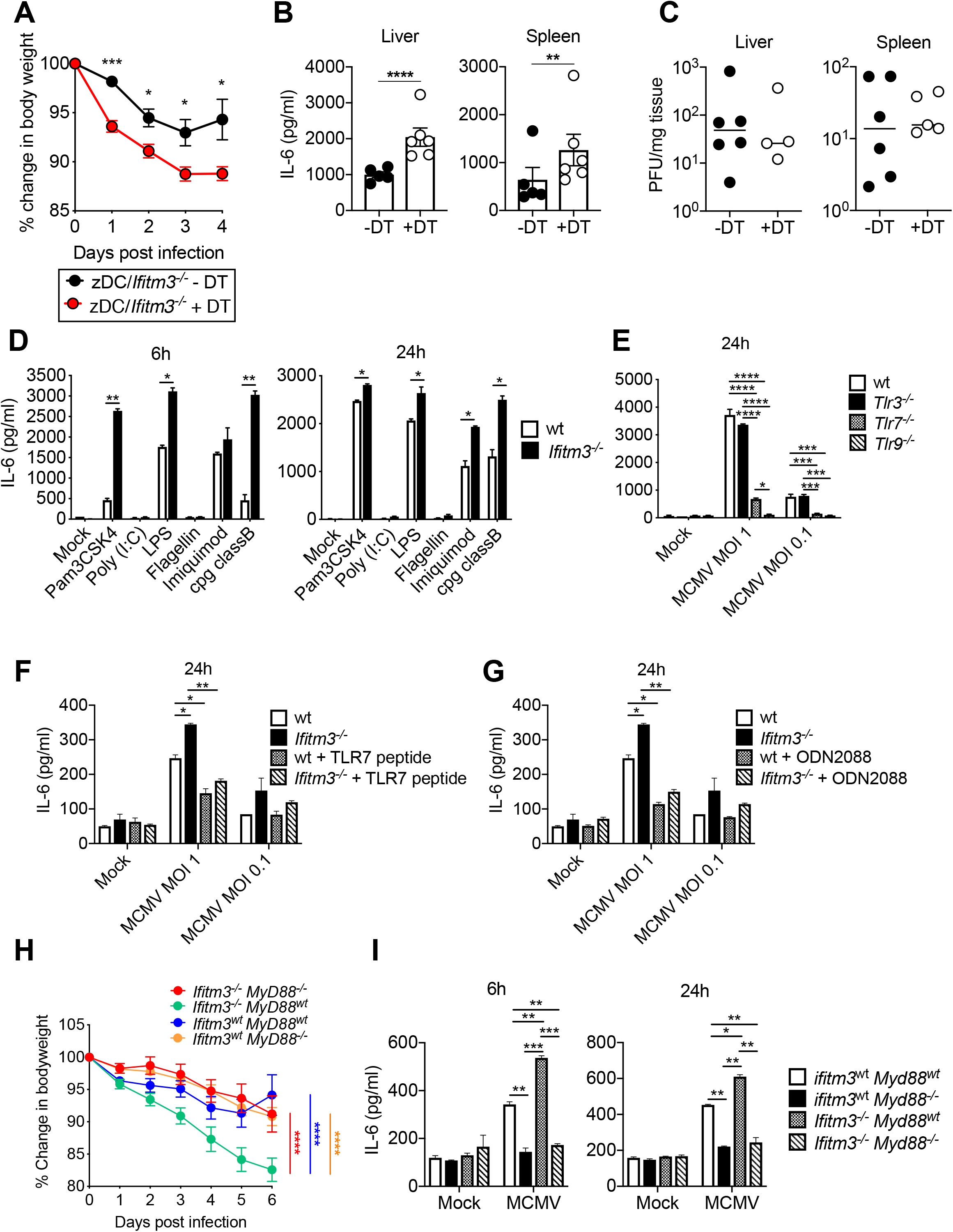
IFITM3 enhances IL-6 downstream of TLRs and MyD88. Mixed bone marrow chimeras with 50:50 of *Ifitm3^-/-^* and wt-zDC-DTR were generated, and treated or not treated with DT. (**A**) Mice were infected with 5 × 10^5^ PFU MCMV and weight loss was assessed over time. (**B**) 4 days p.i. spleens and livers were harvested from MCMV-infected mice, homogenised, and IL-6 was assayed. (**C**) Replicating virus in liver and spleen was quantified 4 days p.i. by plaque assay. Data shown are from 6 (+DT) and 5 (-DT) mice per group. (**D**) BM-DCs from wt and *Ifitm3^-/-^* mice were stimulated with TLR ligands and IL-6 in supernatants was assayed 6h and 24h post stimulation. (**E**) BM-DCs from wt, *Tlr3^-/-^*, *Tlr7^-/-^* and *Tlr9^-/-^* mice were infected with MCMV (MOI 1 or 0.1), and IL-6 in supernatants was assayed 6h and 24h post-infection. (**F**) BM-DCs from wt and *Ifitm3^-/-^* mice were pre-incubated with or without TLR7 synthetic blocking peptide or (**G**) ODN 2088 for 1h prior to infection with MCMV (MOI 1 or 0.1). IL-6 in supernatants was assayed 6h and 24h post infection. (**H**) *Ifitm3^wt^MyD88^wt^*, *Ifitm3^-/-^MyD88^wt^, Ifitm3^wt^MyD88^-/-^, Ifitm3^-/-^MyD88^-/-^* mice were infected with 5 × 10^5^ PFU MCMV and weight loss was assessed over time. Data shown are from 6-8 mice per group. (**I**) BM-DCs from *Ifitm3^wt^MyD88^wt^*, *Ifitm3^-/-^MyD88^wt^, Ifitm3^wt^MyD88^-/-^, Ifitm3^-/-^MyD88^-/-^* mice were infected with MCMV at an MOI of 1, and IL-6 in supernatants was assayed 6h and 24h post infection.

Ifitm3 limits toll like receptor (TLR)-induced cytokine production by bone marrow-derived DCs (BM-DCs, (Stacey et al., 2017)), a result that we reproduced with a broad panel of TLR ligands, with the exception of TLR3 or TLR5 stimulation which induced low IL-6 production in these experiments (Fig. 1D). In addition to cytoplasmic DNA sensing (DeFilippis et al., 2010; Lio et al., 2016), endosomal TLR3, TLR7 and TLR9 are important in the recognition of MCMV and subsequent activation of innate immune responses (Tabeta et al., 2004; Zucchini et al., 2008). Thus, to understand whether endosomal TLRs mediated MCMV-induced IL-6 production in DCs, *Tlr3^-/-^*, *Tlr7^-/-^* and *Tlr9^-/-^* BM-DCs were infected with MCMV and IL-6 was measured (Fig. 1E). *Tlr7^-/-^* and *Tlr9^-/-^* BM-DCs were particularly defective in IL-6 production, suggesting a dominant role for these pattern recognition receptors in MCMV-induced pro-inflammatory cytokine production. In accordance, blocking Tlr7 (Fig. 1F) and Tlr9 (Fig. 1G) signalling significantly reduced IL-6 in both wt and *Ifitm3^-/-^* cells to similar levels following infection with MCMV, suggesting that Ifitm3-mediated regulation of these responses restricts MCMV-induced cytokine production.

The adaptor protein MyD88 is downstream of various TLRs, including TLR7 and TLR9. To provide evidence for a role of TLR signalling in Ifitm3-regulated viral pathogenesis, we crossed *Ifitm3^-/-^* and *Myd88^-/-^* mice and infected them with MCMV. Genetic depletion of MyD88 in *Ifitm3^-/-^* mice resolved infection-induced weight loss to wt levels (Fig. 1H), indicating that signaling through MyD88 is required to drive exacerbated weight loss in *Ifitm3^-/-^* mice. Generation of BM-DCs from *Ifitm3^-/-^/Myd88^-/-^* mice confirmed that IL-6 production post-MCMV exposure was reduced in *Ifitm3^-/-^/Myd88^-/-^* double knockout DCs in comparison to wt *Ifitm3/Myd88* (Fig. 1I). MyD88 also acts downstream of IL-1R (Medzhitov et al., 1998). However, antagonising IL-1R signalling with anakinra has no impact on MCMV weight loss (**Fig. S1A**) or IL-6 production (**Fig. S1B**). Thus, these data suggested that MyD88-dependent TLR signalling drives enhanced pathogenesis in *Ifitm3^-/-^* mice.

### IFITM3 restricts HCMV-induced TLR2-mediated cytokine production by human DCs

Next, we sought to investigate whether IFITM3 regulated virus-induced cytokine production in human DCs. We generated DCs from healthy control (Kolf2) iPSCs, and iPSCs with biallelic mutations in *IFITM3* generated in the Kolf2 background using CRISPR/Cas9 engineering (Wellington et al., 2021). Two clones (*IFITM3^-/-^* F01 and *IFITM3^-/-^* H12) were selected for downstream assays, to control for risk of off-target mutations. Kolf2, *IFITM3^-/-^* H12 and *IFITM3^-/-^* F01 were differentiated into iPS-DCs using previously published methodology (Forbester et al., 2020). DC morphology in culture was similar for all three lines (**Fig. S2A**), and IFITM3 deficiency did not affect the efficiency of DC differentiation, with similar numbers of DC precursors harvested from each line (**Fig. S2B**), and similar surface expression of DC markers CD11c and CD141 (**Fig. S2C**). Furthermore, as *IFITM1* and *IFITM2* have significant sequence homology with *IFITM3*, we determined that mRNA (**Fig. S2D**) and protein (**Fig. S2E**) levels of IFITM1 and IFITM2 were not significantly decreased in iPSCs in comparison to Kolf2 control cells in either *IFITM3^-/^*^-^ clone used in this study. After differentiation into iPS-DCs and challenge with HCMV (merlin strain), IFITM3 protein was detected only in Kolf2 iPS-DCs and not in either *IFITM3^-/-^* line, using a previously validated IFITM3-specific antibody (Makvandi-Nejad et al., 2018; Wellington et al., 2021) (Fig. 2A). In accordance with data generated in the mouse/MCMV system, IL-6 responses were elevated after HCMV stimulation of iPS-DCs that lacked IFITM3 (Fig. 2B). iPS-DCs were non-permissive to productive HCMV infection after low multiplicity of infection (MOI=5), irrespective of IFITM3 expression (**Fig. S3B & S3C**). Thus, IFITM3 suppresses IL-6 production in DCs to both HCMV and MCMV without impacting virus entry.

**Figure 2.**
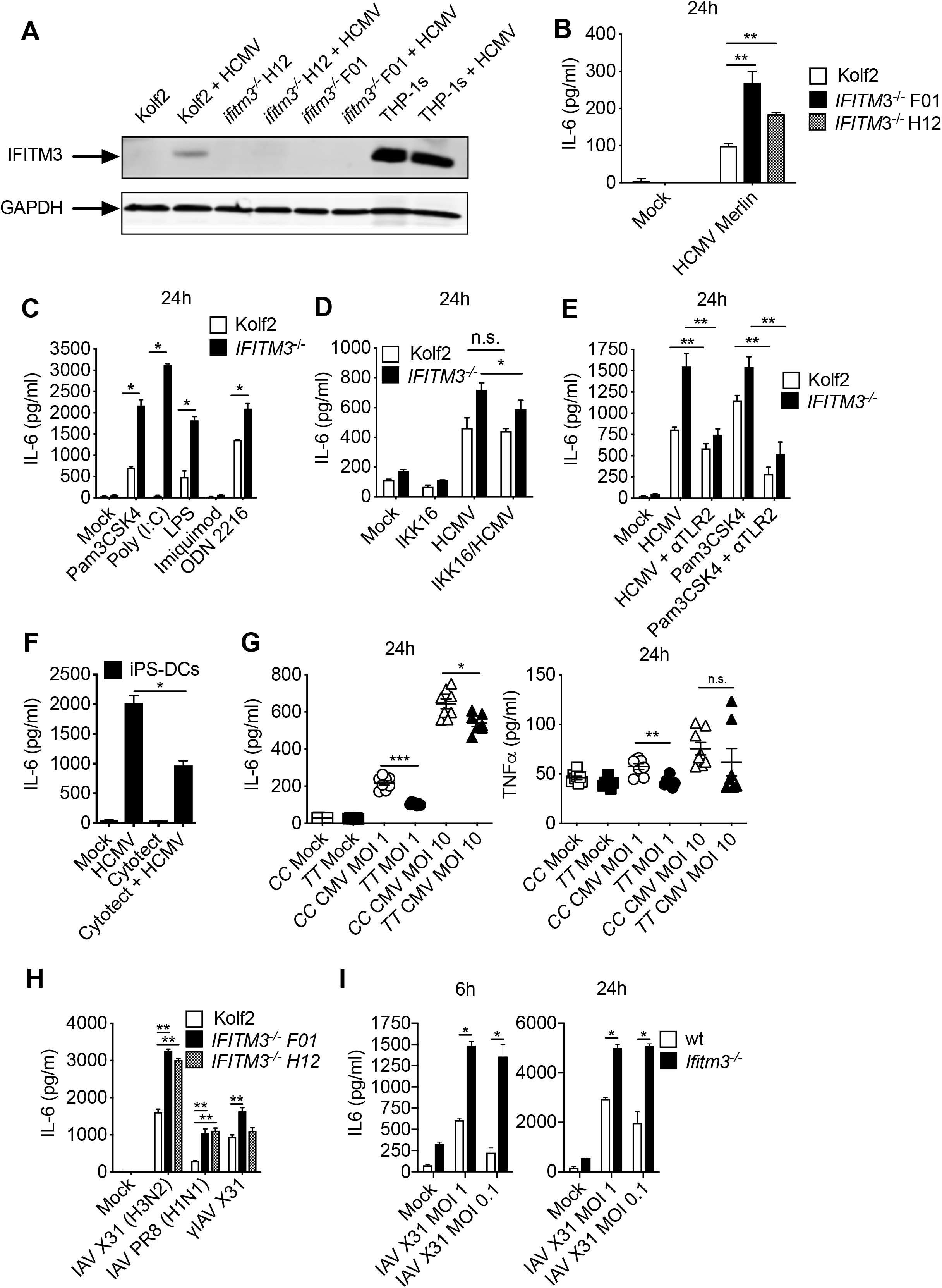
IFITM3 regulates HCMV- and TLR-induced IL-6 in human DCs. ELISA data presented from assays performed in triplicate or more, for at least two independent technical replicates per assay. HCMV (merlin strain) was used for all iPS-DC experiments at MOI 5 unless otherwise stated. (**A**) IFITM3 or GAPDH protein levels after stimulation of control and *IFITM3^-/-^* iPS-DCs for 24h with HCMV were measured by Western blot. Lysate from THP-1 cells was used as a positive control. (**B**) iPS-DCs were stimulated with HCMV, and IL-6 in supernatants were assayed 24h later. (**C**) iPS-DCs were stimulated with TLR ligands and IL-6 was measured. (**D**) iPS-DCs were pre-treated with or without IKK16 for 1h and then stimulated with HCMV and IL-6 measured in supernatants. (**E**) iPS-DCs were pre-treated for 1h with or without neutralizing antibody to TLR2 and stimulated with HCMV or TLR2 ligand Pam3CSK4, and IL-6 was assayed after 24h. (**F**) iPS-DCs were pre-treated with Cytotect CP Biotest, or left untreated, and stimulated with HCMV, with IL-6 in supernatant assayed 24h later. (**G**) Monocyte-derived DCs isolated from human donors genotyped for SNP rs12252 were stimulated with HCMV (MOI 1 or 10), and IL-6 and TNF-α in supernatants were assayed 24h post infection. (**H**) iPS-DCs were stimulated with IAV A/X31 (H3N2), PR8 (H1N1), or gamma-irradiated A/X31 (MOI 1) and IL-6 was measured 6h and 24h later. (**I**) BM-DCs from wt and *Ifitm3^-/-^* mice were infected with IAV A/X31 (H3N2) (MOI 1 or 0.1), and IL-6 in supernatants was assayed 6h and 24h later.

Analogous to data in murine cells, IL-6 secretion was enhanced in IFITM3-deficient cells following stimulation of TLR2, TLR3, TLR4 and TLR9 (Fig. 2C). Antagonizing NFκB using the inhibitor IKK-16, which targets IκB kinase (IKK) (Coldewey et al., 2013), reduced IL-6 in *IFITM3^-/-^* iPS-DCs closer to WT levels, suggesting enhanced IL-6 in IFITM3-deficient cells is partially NFκB dependent (Fig. 2D). This NFκB inhibition had no effect on IL-6 in WT cells, suggesting that in cells expressing IFITM3 IL-6 production is not NFκB-dependent. TLR2 is important for triggering the inflammatory cytokine response to HCMV (Compton et al., 2003) following binding to viral surface glycoproteins gB and gH (Boehme et al., 2006). Neutralizing antibody to TLR2 significantly reduced the IL-6 response that was otherwise amplified in *IFITM3^-/-^* iPS-DCs following exposure to either HCMV or a TLR2 ligand (Fig. 2E). Furthermore, pre-incubation with Cytotect (pooled hyperimmune globulin against HCMV) also reduced IL-6 significantly, providing further evidence for the role of TLR2 in HCMV glycoprotein recognition in mediating subsequent IL6 signalling (Fig. 2F). These data thus suggested that in human DCs TLR2-mediated recognition of HCMV is particularly important for driving the inflammatory response by these cells, and that this inflammatory process was limited by IFITM3. Finally, we generated primary blood monocyte-derived DCs from human donors genotyped for the *IFITM3* SNP rs12252 and stimulated cells with HCMV, with these cells being non-permissive to HCMV infection similarly to iPS-DCs (**Fig. S3D**). Donors with the *CC* allele that associate with reduced IFITM3 function (Zhang et al., 2013) also demonstrated increased IL-6 and TNF-α responses following viral stimulation (Fig. 2G), suggesting that variation within the human *IFITM3* locus could influence differential cytokine responses by myeloid cells in humans.

### IFITM3 regulates DC inflammatory cytokine production in response to evolutionarily diverse pathogenic viruses

*Ifitm3^-/-^* mice are more susceptible to IAV infection, exhibiting loss of viral control, but also alterations in immune responses (Everitt et al., 2012), and variation in human *IFITM3* has been associated with more severe disease after IAV infection (Everitt et al., 2012; Zhang et al., 2020). This suggests that Ifitm3 may regulate pro-inflammatory cytokine responses in response evolutionarily divergent viruses independently of control of virus entry into cells.

We have previously shown in iPS-DCs, IAV-induced IL-6 occurs downstream of sensing by TLR7 and possibly RIG-I (Forbester et al., 2020). Interestingly, when we infected human iPS-DCs with IAV H1N1 (PR8) and H3N2 (X31), we observed no direct IAV restriction (**Fig. S3A**), but significantly enhanced IL-6 was observed in response to infective and γ-irradiated virus (Fig. 2H). In accordance to observations in human DCs, enhanced IL-6 was also observed in murine *Ifitm3^-/-^* BM-DCs in response to IAV (Fig. 2I). Therefore, these data suggest that enhancement of viral induced IL-6 in IFITM3-deficient cells is independent of the viral restriction role IFITM3 plays in other cell types. Thus, overall, our data reveal that IFITM3 regulates NFκB, MyD88 and TLR-dependent virus-induced pro-inflammatory cytokine production and that this is potentially relevant in evolutionarily diverse viruses.

### IFITM3 binds the TLR pathway-associated protein Nogo-B

To determine the mechanism(s) by which Ifitm3 limits TLR-mediated cytokine production, we used a proteomic approach to screen for Ifitm3 binding partners. We performed two separate immunoprecipitation (IP) experiments from cells cultured with differentially labelled amino acids (stable isotype labelling of cells in culture (SILAC)-IP). Both experiments used an Ifitm3-specific antibody to pull down IFITM3, along with any binding partners, from either wt (‘Medium’ labelled) or *Ifitm3*^-/-^ (‘Heavy’ labelled) BM-DCs. In one condition (Fig 3A) cells were mock infected while in the other (Fig 3B) cells were infected with MCMV. Analysis of the ratio of peptides from specific proteins in each condition revealed enrichment of receptor enhancing expression protein 5 (Reep5); Ifitm1 and 2; PRA1 family protein 3 (Arl6ip5, Praf3) and members of the reticulon family of proteins Rtn3 and Rtn4 (Fig. 3A&B).

**Figure 3.**
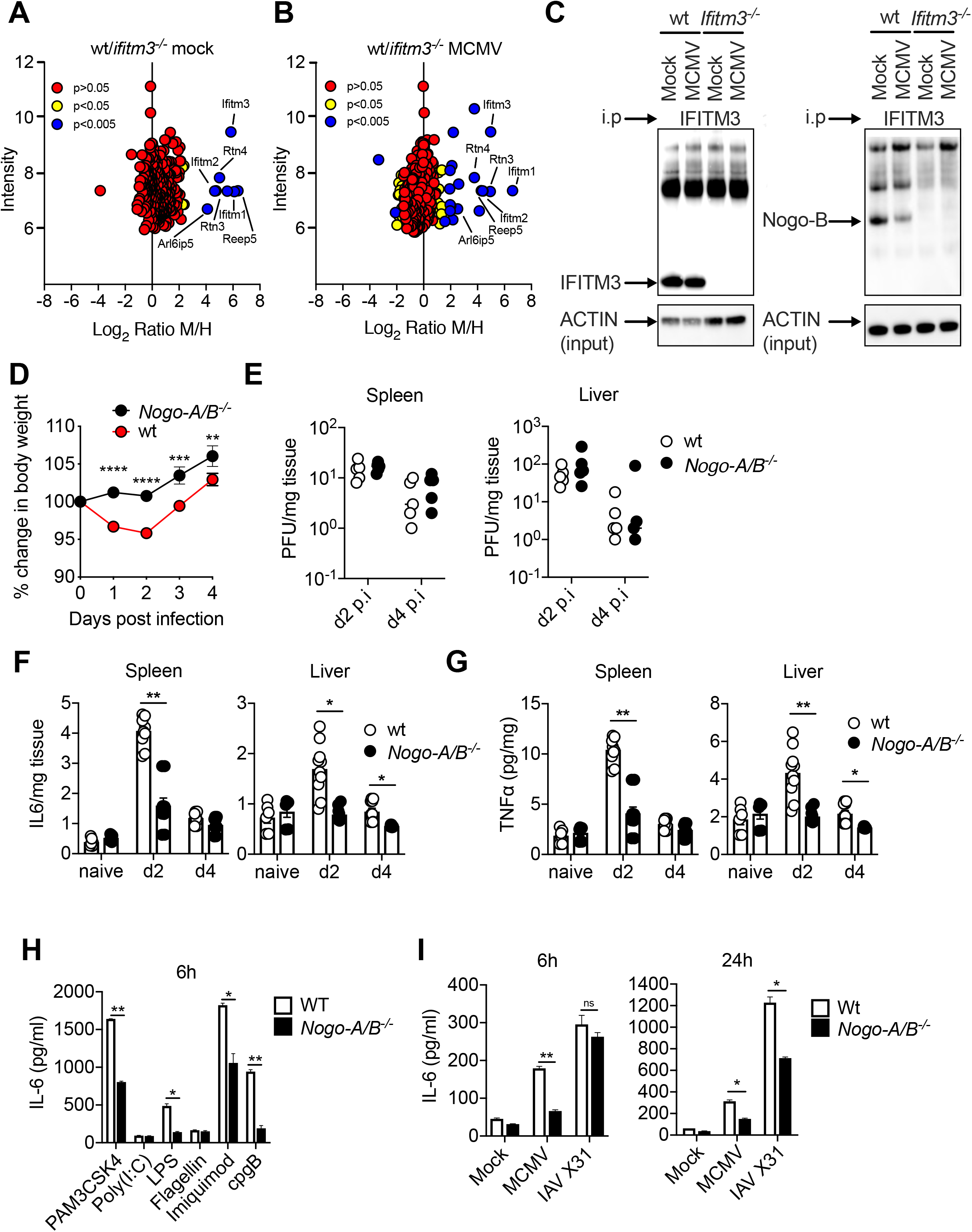
IFITM3 interacts with the reticulon protein Nogo-B. **(A&B)** GM-CSF differentiated BM-DCs from wt and *Ifitm3^-/-^* mice were grown in ‘Medium’ or ‘Heavy’ SILAC medium respectively. Cells were either mock infected (**A**) or infected with MCMV (MOI 1) (**B**) for 3h, lysed and IP for anti-fragilis (IFITM3) was performed. The fold enrichment of each protein is shown. *P* values were estimated using significance A values, then corrected for multiple hypothesis testing (Cox and Mann, 2008). (**C**) BM-DCs from wt and *Ifitm3^-/-^* mice were infected with MCMV (MOI 1) for 3h, lysed and IP for anti-fragilis (IFITM3) was performed. IFITM3, Nogo-B and ACTIN (input samples only) levels were detected by Western blot. (**D**) Wt and *Nogo-A/B^-/-^* mice were infected with MCMV and weight loss was assessed over time. (**E**) Replicating virus from harvested spleens and livers was measured by plaque assay at d2 and d4 p.i. (**F&G**) Harvested spleens and liver tissue supernatant from either naïve, or from d2 and d4 p.i. was assayed for (**F**) IL-6 and (**G**) TNF-α. Data shown represents 5 mice per group from 3 replicate experiments. (**H&I**) BM-DCs from wt and *Nogo-A/B^-/-^* mice were stimulated with or without (**H**) TLR ligands or (**I**) infected with MCMV or IAV A/X31 (MOI 1), for 6h and 24h with IL-6 assayed in supernatants.

Rtn4 encodes several alternatively spliced transcript variants, with the isoforms encoding members of the reticulon family of proteins. The main isoforms are Nogo-A, mainly expressed in the nervous system, Nogo-B, which is ubiquitously expressed in most cell types, and Nogo-C, enriched in skeletal muscle (Oertle and Schwab, 2003). Nogo-B is thought to be involved in maintaining endoplasmic reticulum (ER) shape (Rämö et al., 2016). Intriguingly, however, a regulatory role of Nogo-B in TLR trafficking and expression has been identified (Kimura et al., 2015; Zhu et al., 2017). Interestingly, whereas blotting for Nogo-A of iPS-DC lysates confirmed no Nogo-A expression by myeloid cells (**Fig. S4A)**, we confirmed IFITM-3-Nogo-B interaction in BM-DCs by IP-western blot (Fig. 3C), thus demonstrating that Ifitm3 binds Nogo-B in dendritic cells prior to and following CMV exposure.

### Nogo-B promotes viral pathogenesis

We hypothesized that the interaction between Ifitm3 and Nogo-B was relevant for the immune-regulatory function of Ifitm3. Importantly, MCMV-infected *Nogo-A/B^-/-^* mice exhibited significantly reduced weight loss in comparison to wt mice (Fig. 3D), despite no influence on viral load (Fig. 3E), which was accompanied by reduced pro-inflammatory cytokine production in virus-infected tissue (Fig. 3F&G). Furthermore, IL-6 production by *Nogo-A/B*^-/-^ BM-DCs was significantly reduced following stimulation with TLR2, 4, 7 and 9 ligands (Fig. 3H), and also in response to MCMV and IAV X31 (Fig. 3I). Although *Nogo-A/B^-/-^* mice lack expression of both the Rtn4A and Rtn4B isoforms, we reasoned that ubiquitous expression only of the Nogo-B isoform suggested that only this Rtn4 isoform regulates TLR and virus-induced cytokine production and *in vivo* viral pathogenesis.

We investigated whether Ifitm3 limited viral inflammation by altering Nogo-B abundance. Firstly, we quantified Nogo-B levels post-HCMV exposure in human iPS-DCs and observed a significant enhancement of Nogo-B levels in IFITM3-deficient iPS-DCs at early time-points (1h and, to a lesser extent, 3h, Fig. 4A&B), suggesting that elevated pro-inflammatory cytokine production in IFITM3-deficient cells may reflect elevated Nogo-B activity. In accordance, using siRNAs specific for *IFITM3* and *RTN4* in the monocytic cell line THP-1 in which we could achieve significant knockdown of expression of both proteins (Fig. 4C), we demonstrated that, after exposure to HCMV, Nogo-B knockdown reduced HCMV-induced IL-6 secretion whereas IFITM3 knockdown exacerbated this (Fig. 4D). Importantly, knockdown of both IFITM3 and Nogo-B restored viral-induced IL-6 to level observed with control siRNA (Fig. 4D). We observed similar results in experiments using murine BM-DCs (Fig. 4E&F). These data suggest that in IFITM3-deficient DCs, dysregulated Nogo-B may contribute to enhanced inflammatory responses. However, these experiments were confounded by incomplete Nogo-B (and IFITM3) knockdown by siRNAs. Therefore, to establish complete deletion of Ifitm3 and Nogo-B gene expression, we generated *Ifitm3^-/-^NogoA/B^-/-^* mice and compared MCMV-induced IL-6 secretion by BM-DCs lacking Ifitm3, NogoB or both (Fig. 4G). Importantly, deleting Nogo-B in *Ifitm3^-/-^* BM-DCs reduced virus-induced cytokine production to levels comparable with *NogoA/B^-/-^* BM-DCs and lower than WT cells (Fig. 4G), suggesting a direct association between Ifitm3 and Nogo-B in regulation of virus-induced cytokine production.

**Figure 4.**
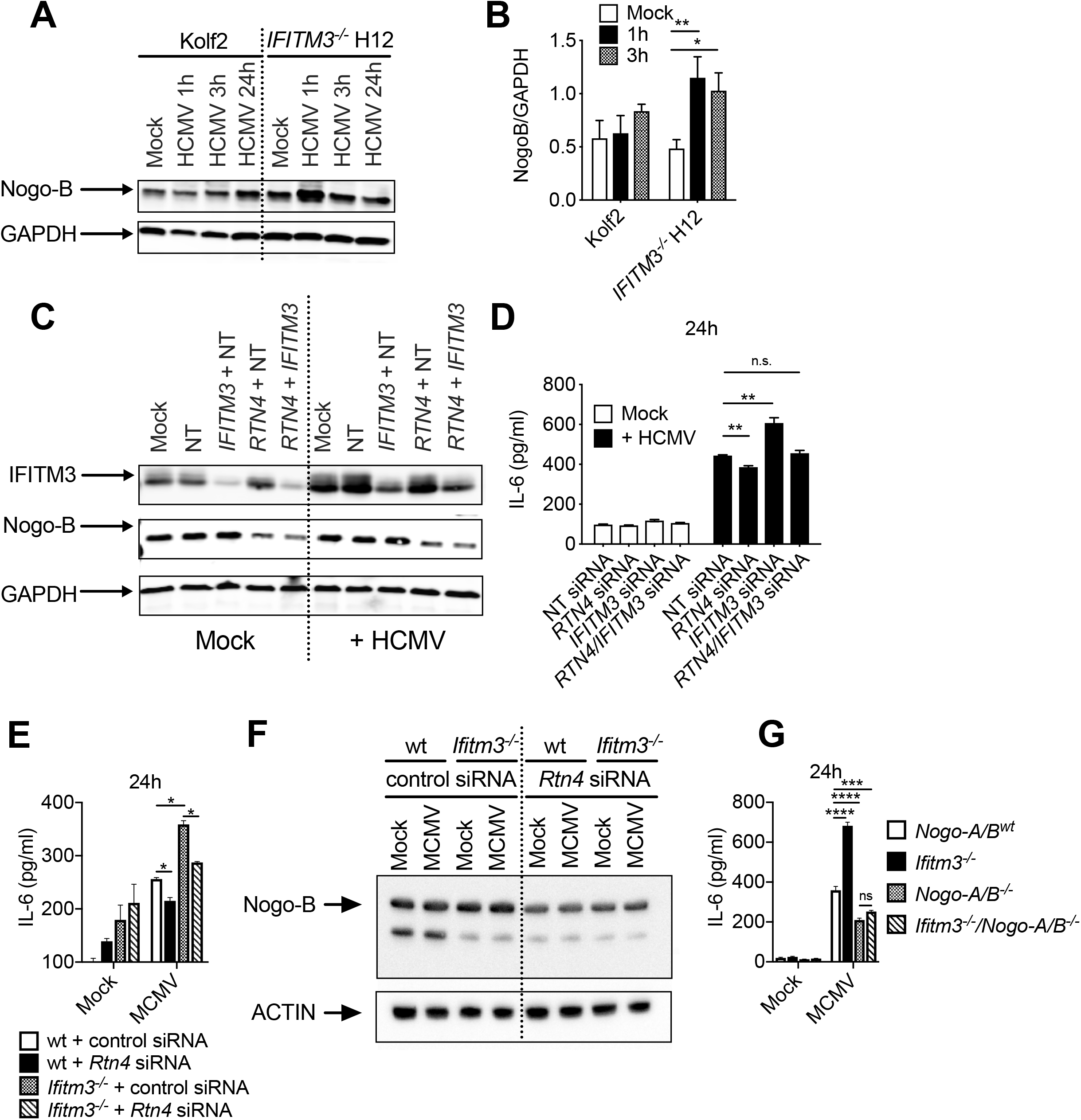
Nogo-B/IFITM3 interaction regulates CMV-induced IL-6. (**A**) Nogo-B and GAPDH was detected by Western blot, after stimulation of Kolf2 and *IFITM3^-/-^* H12 iPS-DCs for 1h, 3h or 24h with HCMV (MOI 5), or mock treatment, and preparation of whole-cell extracts. (**B**) Relative expression of Nogo-B to GAPDH from (**A**) was assessed using ImageJ software. (**C**) THP-1s were treated for 72h with *RTN4*, Non-targeting (NT) and/or *IFITM3* siRNAs and assayed for anti-IFITM3 and anti-Nogo-B by Western blot 72h after siRNA addition and preparation of whole-cell extracts. (**D**) THP-1s treated for 72h with *RTN4*, Non-targeting (NT) and/or *IFITM3* siRNAs were stimulated with HCMV (MOI 5) for 24h, with IL-6 in supernatant measured. (**E**) BM-DCs from wt and *Ifitm3^-/-^* mice targeted with either *Rtn4* targeting siRNA or AllStars control siRNA, and were infected with MCMV (MOI 1), and IL-6 in supernatants was assayed 6h and 24h post infection. (**F**) Nogo-B expression was detected in *Rtn4* siRNA-treated wt and *Ifitm3^-/-^* BM-DCs by Western blot at 3h post-infection with MCMV. (**G**) BM-DCs from *Nogo-A/B^wt^, Ifitm3*^-/-^*, Nogo-A/B^-/-^* and *Ifitm3^-/-^Nogo-A/B^-/-^* mice were infected with MCMV (MOI 1) for 24h with IL-6 assayed in supernatants.

### IFITM3 regulates Nogo-B expression in a proteasomal dependent manner

Nogo-B abundance is regulated via the proteasome (Ahn et al., 2015). In accordance, inhibition of proteasomal cleavage using MG-132 in iPS-DCs (Fig. 5A&B) and murine BM-DCs (where Nogo-B protein was increased in Ifitm3 deficiency, Fig. 5C **& Fig. S1C**) resulted in a significant increase in Nogo-B protein in wt but not IFITM3-deficient cells in both systems. The majority of intracellular proteins are targeted for destruction by the proteasome after tagging with ubiquitin chains (Lecker et al., 2006). In murine BM-DCs generated from wt and *Ifitm3^-/-^* mice, ubiquitin level was measured by western blotting on Nogo-B IP pull-downs, demonstrating a specific reduction of ubiquitination of Nogo-B in both mock and MCMV infected *Ifitm3^-/-^* BM-DCs (Fig. 5E **& Fig. S5A**). Importantly, total ubiquitination in whole cell extracts was not different between wt and *Ifitm3^-/-^* cells (Fig. 5F), suggesting that Ifitm3 specifically affects Nogo-B ubiquitination.

**Figure 5.**
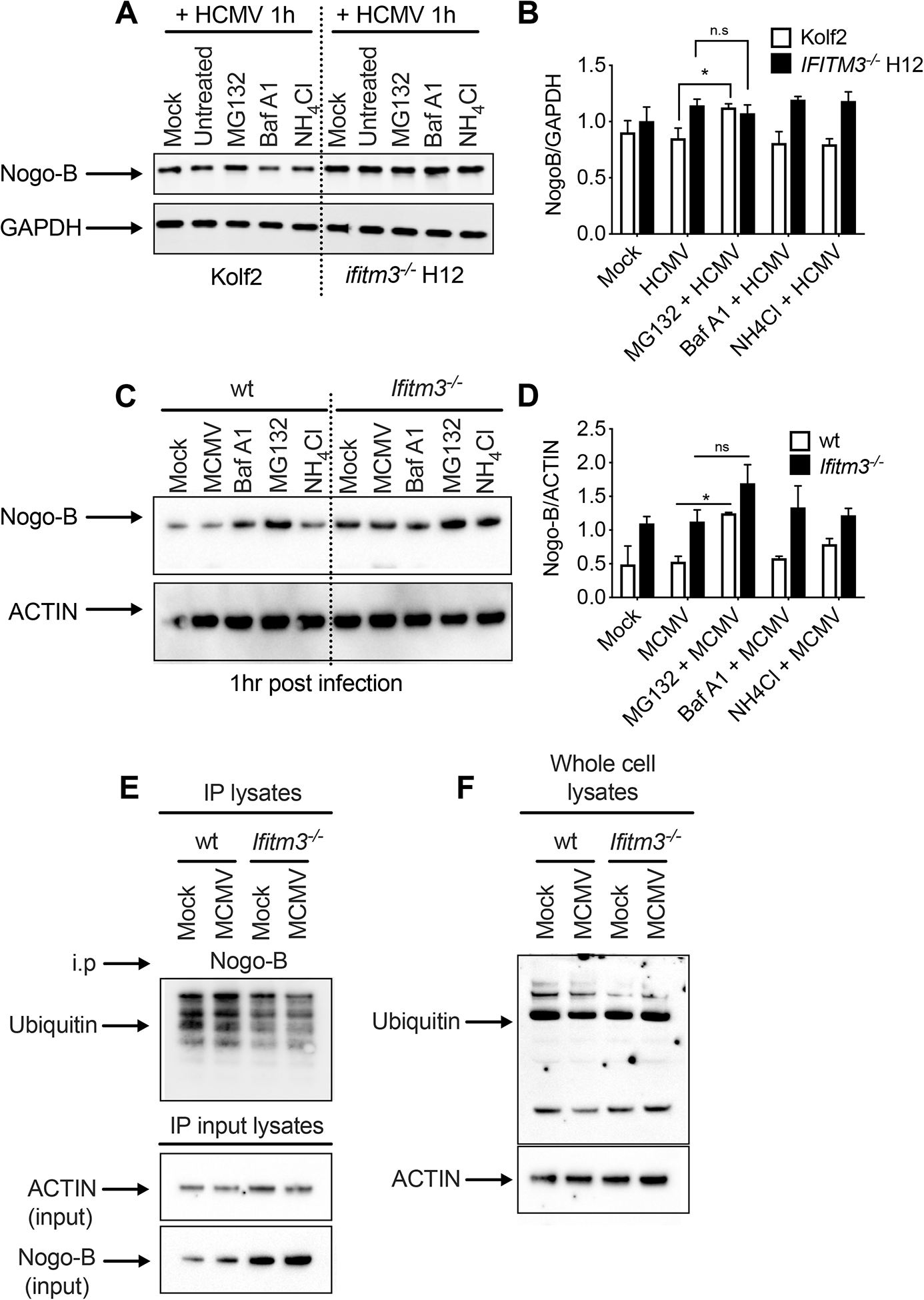
IFITM3 regulates Nogo-B expression via the ubiquitin-proteasome pathway. (**A**) Kolf2, *IFITM3^-/-^* iPS-DCs or (**C**) BM-DCs were pre-treated with Bafilomycin A1, NH_4_Cl or MG132 for 1h, then stimulated with (**A**) HCMV (iPS-DCs) (MOI 5) for 1h or infected with (**C**) MCMV (MOI 1) for 1h (BM-DCs). Nogo-B levels were assayed by Western blot (**A-**iPS-DCs, **C**-BM-DCs), with results from three independent replicates quantified relative to GAPDH (**B**-iPS-DCs) or relative to ACTIN (**D**-BM-DCs). (**E**) BM-DCs from wt and *Ifitm3^-/-^* mice were infected with MCMV (MOI 1) for 1h, lysed and either (**F**) IP for anti-Nogo-B or (**F**) whole cell lysates (no IP) was performed. Anti-Ubiquitin levels of Nogo-B (**E**) and whole cell lysates (**F**) were detected by Western blot. Input samples from (**E**) were also assessed for levels of anti-Nogo-B and equal protein loading using anti-ACTIN via Western blot. (**F**) To verify equal protein loading, the blot was stripped and re-probed with anti-ACTIN antibody.

### IFITM3-Nogo-B interactions regulate TLR2 dynamics

We wanted to understand how IFITM3-Nogo-B interactions influenced virus-induced cytokine production. Given that 1) our data implied a critical role for TLRs in this process and 2) Nogo-B has been implicated in impacting TLR locality within cells (Kimura et al., 2015; Zhu et al., 2017), we examined the impact of IFITM3 deficiency on TLR localization. As TLR2-mediated cytokine responses were regulated by IFITM3 and TLR2 is expressed on the cell surface, we decided to use human cells and TLR2 as a model to investigate whether IFITM3 alters TLR dynamics in response to virus. We performed immunostaining in Kolf2 and *IFITM3^-/-^* H12 iPS-DCs to visualize TLR2 and Nogo-B expression (Fig. 6A). Our observations suggested colocalization (yellow) between TLR2 and Nogo-B after HCMV exposure (Fig. 6A **& Fig. S6A**); and that TLR2 and Nogo-B cellular localization was influenced by IFITM3, with more TLR2/Nogo-B localization visible as cytoplasmic puncta in *IFITM3^-/-^* iPS-DCs 24h post infection (Fig. 6A). The distribution of most Nogo-B protein in these cells did not match a peripheral endoplasmic reticulum pattern. By flow cytometry, we observed significantly more cell surface TLR2 expression by HCMV-exposed Kolf2 iPS-DCs in comparison to mock treated cells (Fig. 6B). In contrast, surface TLR2 expression on *IFITM3^-/-^* DCs does not increase after HCMV exposure and was significantly reduced when compared directly to wt DCs (Fig. 6B). Interestingly, following exposure to the TLR2 ligand Pam3CSK4, surface TLR2 expression reduced rapidly and dramatically in *IFITM3^-/-^* DCs 1h post-Pam3CSK4 exposure, whereas TLR2 surface levels in wildtype Kolf2 cells only reduced later at 4h before rising again by 6h post-exposure (Fig. 6C). We next used our siRNA THP-1 system to assess TLR2 dynamics post-HCMV exposure in response to NogoB/IFITM3 knockdown (Fig. 6D&E). Here, we observed decreased TLR2 surface expression with IFITM3 knockdown at early time-points (1h and 3h) post-virus exposure. In contrast, in cells after Nogo-B knockdown, we observed a less dramatic reduction in surface TLR2 1h and 3h post-HCMV exposure as compared to both non-transfected and IFIMT3 siRNA-treated cells, and markedly increased TLR2 surface expression 24h post-viral exposure. In cells where both Nogo-B and IFITM3 were knocked down, TLR2 surface expression was comparable to control-treated cells (Fig. 6D&E).

**Figure 6.**
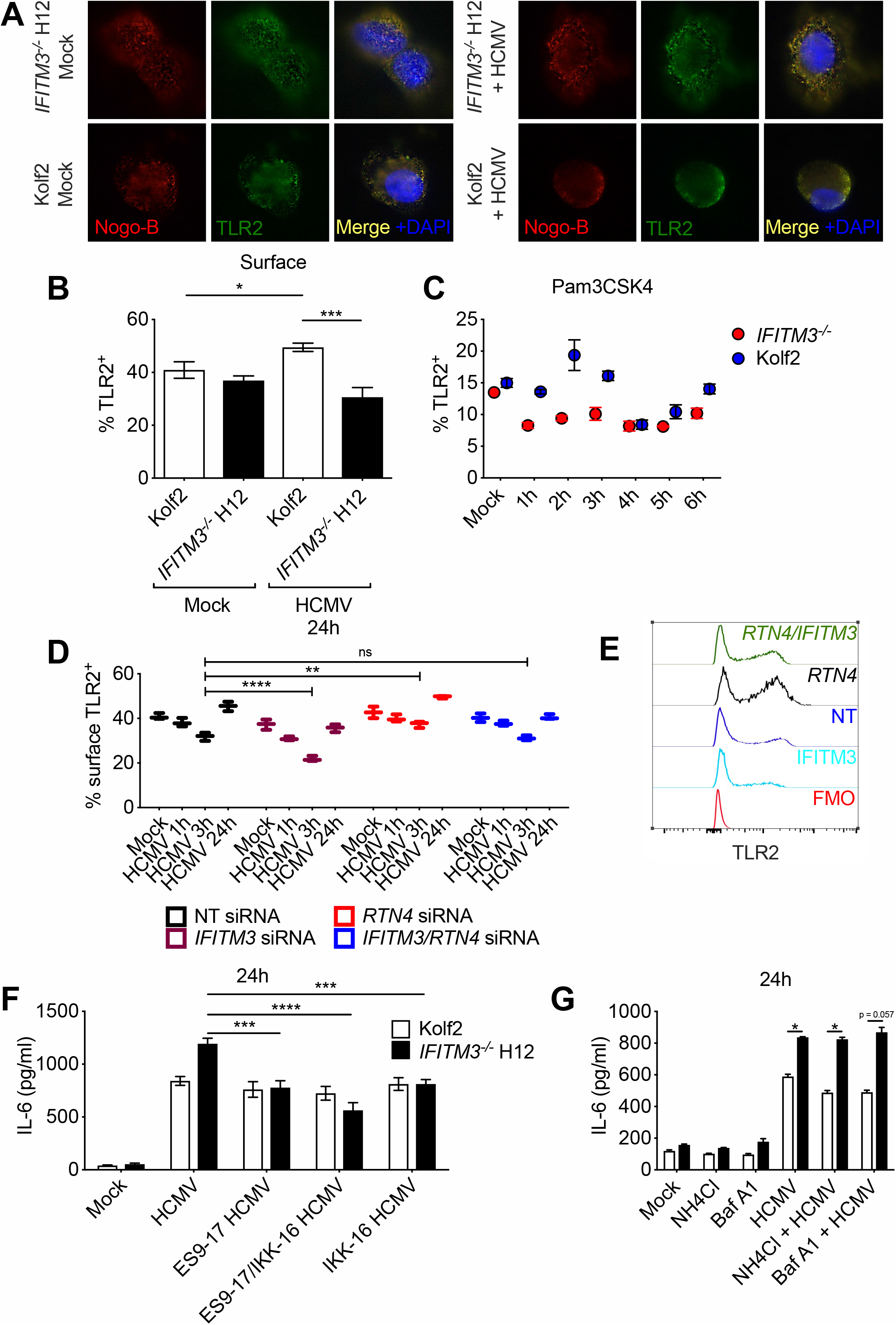
TLR dynamics are altered in IFITM3-deficient DCs. (**A**) Immunostaining for anti-Nogo-B (Red) and anti-TLR2 (Green) in iPS-DCs mock treated or stimulated with HCMV (MOI 5), (Blue; DAPI). (**B**) Surface TLR2 was assessed by flow cytometry in Kolf2 and *IFITM3^-/-^* iPS-DCs either mock treated or stimulated for 24h with HCMV; or (**C**) in iPS-DCs stimulated for 0-6h with TLR2 ligand Pam3CSK4. (**D&E**) Surface TLR2 was assessed by flow cytometry in HCMV-stimulated THP-1s treated for 72h with *RTN4*, Non-targeting (NT) and/or *IFITM3* siRNAs. (**F&G**) IL-6 production by healthy control Kolf2 or *IFITM3^-/-^* H12 iPS-DCs pre-treated for 1 hour with (**F**) endocytosis inhibitor (ES9-17) and/or NFκB inhibitor (IKK-16), or (**G**) Bafilomycin A1 and NH_4_Cl, followed by stimulation with HCMV (MOI 5) for 24h. Data presented is from at least two independent experiments, with samples run in at least triplicate for each ELISA.

Our data imply that Nogo-B regulates TLR2 internalisation and that this process may be regulated by IFITM3. Internalisation of TLR2 and its ligand into endosomes may be important for determining inflammatory cytokines and type I IFN responses, with several studies suggesting that TLR2 generates different signals from different locations within the endosomal pathway, depending on the specific TLR2 ligand used (Barbalat et al., 2009; Brandt et al., 2013; Dietrich et al., 2010; Nilsen et al., 2008). Upon ligand binding, TLR2 is rapidly internalized and trafficked to the Golgi Apparatus (Triantafilou et al., 2006), with evidence in some cell types that signaling from the cell surface could be important for inflammatory cytokine induction whereas endosomal signaling is important for type I IFN induction (Oosenbrug et al., 2017). However, in human monocytes NFκB activation requires, internalisation of TLR2 into endosomal compartments (Brandt et al., 2013). Importantly, when iPS-DCs were treated with the endocytosis inhibitor ES9-17 (Dejonghe et al., 2019), HCMV-induced IL-6 was reduced only in *IFITM3^-/-^* cells (Fig. 6F). Furthermore, concurrent treatment of *IFITM3^-/-^* iPS-DCs with ES9-17 and the NFκB inhibitor IKK-16 had minimal additive effect on HCMV-induced cytokine production, suggesting that in IFITM3-deficient cells enhanced IL-6 is driven by an endocytosis and TLR2-dependent mechanism that is mediated by NFκB signaling. In contrast, treatment with Bafilomycin A1 and ammonium chloride (NH_4_Cl), which inhibit the late stages of autophagy and the acidification of the late endosome, did not impact on IL-6 production in either healthy control Kolf2 or *IFITM3^-/-^* H12 iPS-DCs after HCMV exposure (Fig. 6G). Thus, these data imply that IFITM3 regulates virus-induced inflammatory cytokine production by influencing Nogo-B orchestrated movement of TLR that, in the case of TLR2, is an endocytosis-dependent event.

## Discussion

Herein, we demonstrate using murine and human *in vitro* systems that IFITM3 interactions with the reticulon family member Nogo-B represent a key mechanism for the regulation of inflammatory cytokine production induced by TLRs and evolutionary diverse viruses, and we demonstrate an important role for the Nogo-B/TLR axis in viral pathogenesis *in vivo*. We identified dendritic cells as the key cell type responsible for producing excess IL-6 during MCMV-induced inflammation in IFITM3-deficiency. DCs play a key role in bridging the innate and adaptive immune system (Worbs et al., 2017) and the production of inflammatory cytokines by DCs requires careful balance to ensure sufficient orchestration of antiviral immunity without exacerbating pathology.

Nogo-B, one of the major isoforms of the *RTN4* gene, is a widely expressed reticulon protein that is thought to be involved in formation and stabilisation of the ER (Rämö et al., 2016). However, the role of Nogo-B in immune regulation has been recently suggested, with data indicating a role for Nogo-B in fine-tuning intracellular TLR pathways by influencing their locality within cells, certainly in the case of delivery of TLR9 to endolysosomes (Kimura et al., 2015). Furthermore, Nogo-B promotes surface expression of TLR4 (Zhu et al., 2017)and TLR-induced cytokine production in macrophages (Zhu et al., 2017). In our study, we observed co-localization of TLR2 and Nogo-B, and knockdown of Nogo-B led to alterations in TLR2 movement within the cell after viral stimulation. Thus, it is possible that Nogo-B influenced virus-induced cytokine production by directly altering TLR movement. However, TLR2 is internalized following activation (Triantafilou et al., 2004) and altered TLR2 dynamics in our experiments may reflect alterations in initial activation rather than trafficking per se. Interestingly, Nogo-B regulates the generation of sphingolipids, which are key components of lipid rafts (Cantalupo et al., 2015; Pathak et al., 2018), TLR recruitment to lipid rafts is required for TLR signalling (Triantafilou et al., 2004) and lipid raft disruption reduces inflammatory signalling (Dai et al., 2005; Triantafilou et al., 2002), thus implying that increased Nogo-B activity in IFITM3-deficient cells may lead to an increase in sphingolipid content in lipid rafts and increased inflammatory cytokine production. Interestingly, Nogo-B is also enriched in lipid domains (Acevedo et al., 2004) which may also directly impact on TLR activation. Thus, Nogo-B may alter TLR signalling dynamics by influencing sphingomyelin synthesis and/or lipid raft structure, and this may be relevant for TLRs located either at the cell surface or endosomal compartment.

While reticulon proteins have been implicated in peripheral ER shaping (Voeltz et al., 2006), the data here support an IFITM3-regulated interaction with TLRs at the plasma membrane and in endosomal compartments. Other non-ER examples of plasma membrane and endosomal partners for Rtn proteins include BACE proteinases (He et al., 2004), Cis-prenyltransferases (Harrison et al., 2011; Miao et al., 2006), LILRB2 (Atwal et al., 2008; Huebner et al., 2011) and Rtn4R (Fournier et al., 2001; Laurén et al., 2007). Thus, the regulation of TLR function is consistent with other non-ER Rtn roles. Our immunoprecipitation studies revealed the presence of both Rtn3 and Rtn4 in IFITM3 complexes. While functional studies of Rtn4 deletion yielded prominent effects on CMV induced cytokine production, the role of Rtn3 may be partially redundant and combined Rtn3/4 deletion may yield even greater effect. While the current study focuses on viral infection and dendritic cells, the prominent expression of Rtn’s in the brain suggests that the mechanisms defined here may contribute in the regulation of TLR function for a range of neuro-inflammatory diseases.

The exact mechanisms through which Ifitm3 promotes ubiquitination and how this Nogo-B-Ifitm3 axis influences TLR signalling remains to be determined. However, it is clear from our studies that in IFITM3-deficient cells where we observed increased Nogo-B expression, we also observed heightened inflammatory cytokine production and internalisation of TLR2. It was initially thought that TLRs were partitioned into surface or endosomal TLRs, with all signalling occurring in either location. However more recent studies have demonstrated activation of different inflammatory mediators, dependent on differential TLR locality within the cell (Kagan et al., 2008; Sasai et al., 2010). Interestingly, internalisation of cell-surface TLR2 into endosomal compartments in human monocytes is required for NFκB activation (Brandt et al., 2013), and TLR2-induced IL-6 expression can be restricted by endocytosis inhibition (Petnicki-Ocwieja et al., 2015). Our observation that inhibition of endocytosis or NFκB induces a similar reduction of IL-6 production only in IFITM3-deficient cells implies that a similar process may be suppressed in IFITM3-expressing DCs to reduce virus-induced inflammation.

Interestingly, the work of Kimura *et al* that revealed a role for Nogo-B in regulation of TLR locality in macrophages reported no impact of Nogo-B on TLR2-induced cytokine production (Kimura et al., 2015), in contrast to our data. These differences may reflect the different cell types studies. Indeed, signalling pathways downstream of TLR2 have previously been observed to be divergent in DCs in macrophages (Groft et al., 2020), implying differential regulation of TLR-induced cytokine responses in different myeloid cells. In support of this hypothesis, Ifitm3 has no impact on MCMV-induced cytokine responses by murine macrophages (Stacey et al., 2017).

In our study, we used iPS-DCs as a model for human myeloid cells. These cells provide a useful model for studying human cells *in vitro*, due to their unlimited numbers, and the ability to genetically edit the iPSC progenitors. In these cells, IFITM3 did not directly restrict IAV entry, in contrast to other cell types (Brass et al., 2009). It has been shown that CD141^+^ human DCs are resistant to productive infection by IAV (Silvin et al., 2017). Here, we measured viral NP expression, which is present in viral particles, suggesting that uptake of virus is not affected by IFITM3. Similarly, the relatively low MOI for HCMV used in our studies also led to no productive infection despite significant virus-induced cytokine production. Therefore, in both human experimental systems we were able to disentangle the immune-regulatory and antiviral functions of IFITM3 in response to evolutionary diverse viruses.

For many viral diseases, effective treatments remain elusive, and vaccines are not readily available or quickly distributed in a pandemic scenario. Therefore, immunomodulatory approaches to reduce host inflammation during viral infection can provide attractive alternatives for rapid treatment of infected individuals, as highlighted by the SARS-CoV-2 pandemic (Abani et al., 2021). Our study reveals Nogo-B as a novel interacting binding partner of IFITM3 and highlights the previously unappreciated role for both IFITM3 and Nogo-B in influencing virus-induced inflammatory events. Targeting this interaction and dissecting the downstream impacts of this on innate immune recognition may help to expand strategies for controlling pathologic viral induced-inflammation.

## Supporting information

Supplemental RAW proteomics data

## Supplemental Figures

**Figure S1. Antagonising IL-1R signalling has no impact on MCMV-induced weight loss or cytokine production.** (**A**) Wt and *Ifitm3^-/-^* mice were infected with MCMV with either 25mg/kg Anakinra or PBS and weight loss was assessed over time. Data shown represents 4 mice per group. (**B&C)** BM-DCs from wt and *Ifitm3^-/-^* mice were pre-incubated with or without (**B**) Anakinra and then infected with MCMV (MOI 1). Supernatant was harvested at 6h and 24 post infection and IL-6 levels were assayed.

**Figure S2. IFITM3-deficiency does not affect DC differentiation and phenotype.** CRISPR/Cas9 was used to generate biallelic mutations in IFITM3 in Kolf2 iPSCs. iPSCs were differentiated into dendritic cells using defined concentrations of growth factors to generate embryoid bodies (EBs), GM-CSF and IL-4 to generate immature DCs from Ebs. (**A**) Morphology of iPS-DCs in culture (100X). (**B**) Total cell numbers of DC precursors harvested from DC differentiation plates. Data shown are from 4 independent differentiations per iPSC line. (**C**) Surface expression of DC markers CD11c and CD141 was examined by flow cytometry. Representative plots presented from one experiment, with experiments performed at least three times. (**D**) Gene expression of *IFITM1* and *IFITM2* by iPS-DCs relative to *GAPDH* was quantified using TaqMan gene expression assays. Data presented shows mean Ct values from four technical replicates, with assays repeated in triplicate. (**E**) IFITM1 and IFITM2 expression in iPSCs was assayed by Western blot.

**Figure S3. IFITM3-deficiency in iPS-DCs does not alter infectivity by IAV or HCMV**. (**A**) *IFITM3^-/-^* and Kolf2 iPS-DCs were stimulated with IAV A/X31 (MOI 1) and then stained for IAV NP 24h post-stimulation. (**B&C**) *IFITM3^-/-^* and Kolf2 iPS-DCs were non-permissive to HCMV using both merlin and TB40, as assayed by expression of IE2-GFP, with representative FACS plots from these assays, and a permissive cell line (macrophages), shown in (**B**). (**D)** Monocyte-derived DCs were non-permissive to HCMV, as assayed by expression of IE2-GFP, with representative FACs plots shown for *CC* or *TT* donors.

**Figure S4. iPS-DCs do not express NOGO-A**. Expression of anti-Nogo-A was assayed in iPS-DCs, A549s and HepG2s from whole cell lysates by Western blot.

**Figure S5. Ifitm3 specifically affects Nogo-B ubiquitination.** (**A**) wt and *Ifitm3^-/-^* BM-DCs were infected with MCMV (MOI 1) for 3h. Post infection, IP with anti-Nogo-B or anti-sheep IgG control antibody was performed, and whole cell lysates were generated. Expression of Nogo-B (top) was measured via Western blot on IP lysates whereas IP input controls were measured for expression of ACTIN (bottom).

**Figure S6.** (**A**) **Nogo-B and TLR2 colocalise after HCMV infection.** (**A**) Immunostaining for TLR2 and NogoB was performed in *IFITM3^-/-^* and Kolf2 iPS-DCs stimulated for 1h, 3h or 24h with HCMV or mock treated followed by colocalization analysis. Data presented is percentage colocalization for at least 10 cells per condition.

**Supplemental file – SILAC_proteingroups**. Full dataset of SILAC IP analysis.

## Methods

### Mice, viral infections and treatments

IFITM3-deficient (*Ifitm3^-/-^*) and wt control mice have been described previously (Everitt et al., 2012) and were crossed with *Myd88^-/-^* mice for some experiments. Age- and sex-matched mice between 7 and 12 weeks of age were used in the experiments. *Tlr3^-/-^*, *Tlr7^-/-^*and *Tlr9^-/-^* mice were a kind gift from Caetano Reis e Sousa (Crick Institute, London). MCMV (pSM3fr-MCK-2fl BACmid) was grown and titred using 3T3 cells with a carboxycellulose overlay. Mice were infected via intraperitoneal (i.p.) injection with between 5 × 10^5^ to 2 × 10^6^ PFU MCMV. For Anakinra treatment, mice were injected i.p. with Anakinra (KINERET: Cardiff & Vale NHS Pharmacy) (25mg/kg) or PBS control on day 0 p.i. For infectious virus quantification form harvested tissue, viral load was determined via plaque assay as previously described (Stack et al., 2015).

### HCMV preparation for human in vitro assays

HCMV strain merlin (pentamer deficient) or TB40-BAC4 was propagated in human foreskin fibroblasts (HFF)-TERTs as previously described (merlin – (Stanton et al., 2010)), (TB40-BAC4 – (Sinzger et al., 2008)) with viral PFU calculated using a previously published method (Stanton et al., 2010).

### Generation of z-DC/*Ifitm3^-/-^* Chimeric mice

Wt recipient mice were gamma irradiated with 550rad for 2min over two intervals. After 24h, mice were intravenously (i.v) transfused with 1 × 10^6^ bone marrow derived cells from *Ifitm3^-/-^* and wt-zDC-DTR (Dimonte et al., 2021) donors. 12 weeks later mice were administered either with or without Diptheria toxin (DT) (1μg/ml) intraperitoneally (i.p). 24h later mice were infected with MCMV and weight loss was assessed over time.

### *In vitro* infections

Bone marrow derived dendritic cells (BM-DCs) cells from either wt, *ifitm3^-/-^, Tlr3^-/-^*, *Tlr7^-/-^* and *Tlr9^-/-^, Nogo-A/B^-/-^, Nogo-A/B^wt^, Ifitm3^wt^MyD88^wt^*, *Ifitm3^-/-^MyD88^wt^, Ifitm3^wt^MyD88^-/-^, Ifitm3^-/-^MyD88^-/-^ and Ifitm3^-/-^Nogo-A/B^-/-^* mice were incubated at 4 × 10^5^ cells per ml in R10 media supplemented with 100U/mL Penicillin/streptomycin, 2mM L-glutamine, 0.1M HEPES, 50μM B-Mercaptoethanol and 1 × MEM non-essential amino acids (all Gibco, Thermo-Fisher) and 20ng/ml GM-CSF (Biolegend) for 9 days replenishing the media at d2, 4 and 7. Differentiated BM-DCs were then stimulated with MCMV at a multiplicity of infection (MOI) of 1 or 0.1 or IAV A/X-31 (H3N2) at an MOI of 1 as indicated. Cells were infected via centrifugation and supernatant was harvested at 6h and 24h post infection. In some experiments, cells were treated with various blocking reagents, TLR ligands or siRNAs as described in more detail below.

### iPSC line generation and culture

The healthy control human iPSC line Kolf2 was acquired through the Human Induced Pluripotent Stem Cells Initiative Consortium (HipSci; www.hipsci.org), through which it was also characterized (Leha et al., 2016). Consent was obtained for the use of cell lines for the HipSci project from healthy volunteers. Prior to differentiation, iPSCs were grown feeder-free using the Essential 8 Flex Medium kit (Thermo-Fisher) on Vitronectin (VTN-N, Thermo-Fisher) coated plates as per manufacturer’s instructions to 70-80% confluency. iPSCs were harvested for differentiation using Versene solution (Thermo-Fisher).

### Generation of *IFITM3^-/-^* iPSCs

The Wellcome Trust Sanger Institute core gene-editing pipeline generated *IFITM3^-/-^* iPSC lines, as previously described in (Wellington et al., 2021). The knockout of IFITM3_F01 was generated by a single T base insertion in the first exon using CRISPR/Cas9 in the Kolf2_C1 human iPSC line (a clonal derivative of Kolf2). This was achieved by nucleofection of 10^6^ cells with Cas9**<u>-</u>**crRNA-tracrRNA ribonucleoprotein (RNP) complexes. Synthetic RNA oligonucleotides (target site: 5’-TGGGGCCATACGCACCTTCA CGG, WGE CRISPR ID: 1077000641, 225pmol crRNA/tracrRNA) were annealed by heating to 95°C for 2min in duplex buffer (IDT) and cooling slowly, followed by addition of 122pmol recombinant eSpCas9_1.1 protein (in 10mM Tris-HCl, pH 7.4, 300mM NaCl, 0.1mM EDTA, 1mM DTT). Complexes were incubated at room temperature for 20min before electroporation. After recovery, cells were plated at single cell density and colonies were picked into 96 well plates. 96 clones were screened for heterozygous and homozygous mutations by high throughput sequencing of amplicons spanning the target site using an Illumina MiSeq instrument. Final cell lines were further validated by Illumina MiSeq. Two homozygous targeted clones were used in downstream differentiation assays.

IFITM3_WT sequence: CCTCTGAGCATTCCC<u>TGGGGCCATACGCACCTTCA</u>CGGAGTAGGCGA

IFITM3_F01 MUT sequence: CCTCTGAGCATTCCC<u>TGGGGCCATACGCACCTTTCA</u>CGGAGTAGGCGA

Insertion of T

IFITM3_H12 First Allele MUT sequence: CCTCTGAGCATTCCC<u>TGGGGCCATACGCACCTTTCA</u>CGGAGTAGGCGA

Insertion of T

IFITM3_H12 Second Allele MUT sequence: CCTCTGAGCATTCCC<u>TGGGGCCATACGCACCTCTTCA</u>CGGAGTAGGCG

Insertion of CT

WT Protein: MNHTVQTFFSPVNSGQPPNYE<u>M</u>LKEEHEVAVLGAPHNPAPPTSTVIHIRSETSVP<u>DHVVWSLFNTLFMNPCCLGFIAFAYSVKSRDRKMVGDVTGAQAYASTAKCLNIWALILGILMTIL</u>LIVIPVLIFQAYG*

MUT Protein (T Insertion): MNHTVQTFFSPVNSGQPPNYE<u>M</u>LKEEHEVAVLGAPHNPAPPTSTVIHIRSETSVP<u>DHVVWSLFNTLFMNPCCLGFIAFAYSVK</u>V*

MUT Protein (CT Insertion): MNHTVQTFFSPVNSGQPPNYE<u>M</u>LKEEHEVAVLGAPHNPAPPTSTVIHIRSETSVP<u>DHVVWSLFNTLFMNPCCLGFIAFAYSVK</u>SLGTGRWLAT*

Underlined sequences are CD225 domain, and alternative methionine start codon in some protein isoforms.

### Differentiation of iPSCs to iPS-DCs

Differentiation of iPSCs to dendritic cells and macrophages. To differentiate iPSCs to dendritic cells, slight modifications were made to a previously published protocol (Sachamitr et al., 2018). Briefly, upon reaching confluence, iPSCs were harvested and plated into Essential 8 Flex medium supplemented with 50ng/ml bone morphogenetic protein 4 (BMP-4; Bio-Techne), 20ng/ml stem cell factor (SCF; Bio-Techne), 50ng/ml vascular endothelial growth factor (VEGF; Peprotech EC Ltd.), and 50ng/ml GM-CSF (Peprotech EC Ltd.) in ultralow attachment (ULA) plates (Corning). The medium was changed to X-VIVO-15 (Lonza), with sequential removal of BMP-4 by day 5, VEGF by approximately day 14, and SCF by approximately day 19. In addition, IL-4 (Peprotech EC Ltd.) was added sequentially in increasing concentrations, starting from approximately day 12 at 25ng/ml and increasing to 100ng/ml by approximately day 20. By day 20, floating immature DCs were harvested from ULA plates, filtered through 70-μm filters (Corning), counted, and seeded at 1 × 10^6^ per well of 6-well CellBind plates (Corning) in X-VIVO-15 medium supplemented with 100ng/ml IL-4 and 50ng/ml GM-CSF. iPS-DCs were used for assays at the immature phase between 4 and 5 days postseeding in CellBind plates. For the assays, floating iPS-DCs were harvested from differentiation plates, washed with PBS, counted, and seeded in X-VIVO-15 medium without cytokines at an assay-dependent concentration.

### Genotyping of human donors

*IFITM3* rs12252 genotype was identified for each participant through PCR amplification of the *IFITM3* gene using primers: Forward ggcagaggtgagggcttt, Reverse gtcccttacgagtctcccac. 100ng of genomic DNA was added to Taq DNA polymerase (PCR Biosystems) and samples were run at: 95°C 15 seconds, 58°C 15 seconds, 72°C 15 seconds for 35 cycles. Amplification was confirmed by running samples on a 2% agarose gel prior to PCR Clean up (QiaQuick PCR Purification kit, Qiagen). Samples were Sanger sequenced (Source Bioscience) and rs12252 was identified from the DNA Electropherogram File (.ab1 file).

### Generation of blood-derived human dendritic cells

PBMCs from three independent donors were isolated from leukapheresis products using Lymphoprep density gradient centrifugation and SepMate PBMC isolation tubes (StemCell Technologies), under the Weatherall Institute of Molecular Medicine, University of Oxford Human Tissue Authority license 12433. Human CD14 microbeads were used in combination with LS columns (both Miltenyi Biotec) to positively select CD14^+^ blood monocytes. CD14^+^ cells were seeded at a density of 3 × 10^6^ to 5 × 10^6^ isolated monocytes in 3ml of RPMI medium supplemented with 10% heat-inactivated fetal bovine serum (FBS; Sigma-Aldrich), 250IU/ml IL-4, and 800IU/ml GM-CSF (both Peprotech EC Ltd) into a 6-well plate and incubated at 37°C for 2 days. After 2 days, 1.5ml of medium was removed from each well, and 1. ml of fresh medium supplemented with 500IU/ml IL-4 and 1,600IU/ml GM-CSF was added. After a further 3-day incubation, cells were harvested at the immature phenotype and assayed.

### Stimulation of human DCs with IAV and HCMV

iPS-DCs, or human monocyte-derived DCs (mDCs) were stimulated with A/X-31 influenza virus or gamma-irradiated A/X-31 influenza virus at MOI of 1, or HCMV strain merlin at MOIs stated in each legend, by the addition of virus to a small volume of X-VIVO-15 cell culture supernatant (50uL for assays in 96 well plates; 200uL for assays in 24 well plates) followed by incubation at 37°C for 1h, after which fresh culture medium was added. After 6h and 24h inoculation respectively, 150ul supernatant was collected and stored at −80°C. Each infection condition was repeated in triplicate.

### Inhibitor assays

For proteasomal/lysosomal inhibition assays, 5 × 10^5^ cells per well iPS-DCs (human) or 1 × 10^5^ BM-DCs (mouse) per condition were pre-incubated for 1h prior to viral stimulation or infection with inhibitors MG132 (10μM; Merck Millipore) BafA1 (0.5μM; Invivogen) or NH_4_Cl (10mM; Sigma-Aldrich), before addition of HCMV for 1h and 3h or with MCMV for 6h and 24h as described above, after which DCs were harvested for protein preparation or supernatants harvested. Further inhibition assays were performed using 1 × 10^5^ cells per well BM-DCs (mouse) per condition. Cells were pre-incubated for 1h prior to viral infection with either TLR7 synthetic peptide (2μg/ml; Thermo-Fisher), ODN 2088 (10μM; Invivogen), 3-Methyladenine (3MA, 5mM; Merck Millipore) or Anakinra (500ng/ml; Cardiff & Vale NHS Pharmacy) with either protein lysates generated or supernatants harvested. For human blocking assays that target endocytosis and NFκB signalling, ES9-17 (endocytosis inhibitor; 100μM; Merck Millipore) and/or IKK-16 (NFκB inhibitor; IKK inhibitor VII, 1nM; Cambridge Bioscience) were added 1h prior to addition of HCMV. Supernatants were harvested after 24h. For assays using neutralising antibodies to TLR2 (200μg/mL; Invivogen) or HCMV (500μg/ml; Cytotect CP Biotest) antibodies were added 1h prior to addition of HCMV or TLR2 ligand.

### Preparation of RNA and RT-qPCR

iPS-DCs were harvested from the plates, and RNA was prepared using the RNeasy minikit (Qiagen). RNA was reverse transcribed with the QuantiTect reverse transcription (RT) kit (Qiagen), according to the manufacturer’s protocol. All RT-qPCR experiments were performed with TaqMan gene expression assays and TaqMan gene expression master mix (Applied Biosystems) on the Applied Biosystems StepOne real-time PCR system. RT-qPCR data were analyzed via the comparative threshold cycle (*C_T_*) method with glyceraldehyde 3-phosphate dehydrogenase (GAPDH) as an endogenous control.

### TLR ligand stimulation

Human iPS-DCs were plated at 2 × 10^4^ cells per well in 200μl of X-VIVO-15 medium without cytokines. TLR ligands were added directly to the medium, and supernatants were harvested after a 24h incubation at 37°C. For the assays, TLR ligands were used at the following concentrations: TLR2; Pam3CSK4, 300ng/ml (InvivoGen); TLR3; Poly(I:C), 50μg/ml (InvivoGen); TLR4; Lipopolysaccharide (LPS), 500ng/ml (Sigma-Aldrich); TLR7; Imiquimod, 50μg/ml (InvivoGen); and TLR9; ODN 2216, 3μg/ml (Miltenyi Biotech). Mouse BM-DCs were plated out at 1 × 10^5^ cells per well. TLR ligands were added directly to the well as before with supernatants harvested at 6h and 24h at 37°C. TLR ligands were used at the following concentrations: TLR2; Pam3CSK4, 0.5μg/ml (InvivoGen); TLR3; Poly(I:C), 10μg/ml (InvivoGen); TLR4; LPS, 10μg/ml (InvivoGen); TLR5; Flagellin 5μg/ml (InvivoGen); TLR7; Imiquimod, 5μg/ml (InvivoGen); and TLR9; CpG Class B ODN 1826, 0.05μM (Invivogen).

### Cytokine analysis

Human and mouse IL-6 and TNF-α protein were measured by an enzyme-linked immunosorbent assay (ELISA) (BioLegend) according to the manufacturer’s instructions.

### Proteomic pulldowns

Bone marrow derived dendritic cells (BM-DCs) cells from either wt or *ifitm3*^-/-^ mice were grown in SILAC RPMI media (Gibco, Thermo-Fisher) either supplemented with 10% HI dialysed and filtered FCS (Sigma-Aldrich) 0.1M HEPES, 50μM B-Mercaptoethanol (both Gibco, Thermo-Fisher), and either L-Lysine-2HCl 13C6 15N2 and L-Arginine-HCl 13C6 15N4 (‘Heavy’ amino acids) or L-Lysine-2HCl 4,4,5,5-D4 and L-Arginine-HCl 13C6 (‘Medium’ amino acids) (all Cambridge Isotope Laboratories). Wt cells were grown in ‘Medium’ SILAC media and *ifitm3*^-/-^ were grown in ‘Heavy’ SILAC media. BM-DC cells were differentiated as described above for 10 days and were infected or not with MCMV at an MOI of 1 for 3h. Cells were removed from the plates post infection and subsequently lysed using Pierce™ IP lysis buffer (Thermo-Fisher) supplemented with 1M proteasome inhibitors (Sigma-Aldrich). Immuno-precipitation (IP), for IFITM3 (α-fragillis, 1μg/ml; Abcam) was performed on all samples as described previously using Pierce™ Protein Plus Agarose A/G beads (Thermo-Fisher) (Stanton et al., 2014). To confirm specificity of IP, control anti-rabbit IgG (1μg/ml; Abcam) was also performed. Post IP beads were that were bound to IFITM3 only were combined and eluted from the Agarose using 1 × NuPAGE^TM^ LDS sample buffer (Thermo-Fisher) supplemented with 100mM DTT (Sigma-Aldrich). Samples were run on a NuPAGE^TM^ 4 to 12% Bis/Tris gels (Thermo-Fisher) running approximately 0.5 to 1cm into the gel. The gel was stained using Colloidal blue staining kit (Thermo-Fisher) as per manufacturers recommendations. The stained lane was excised and cut into 6 fragments. Following in-gel reduction and alkylation, proteins were digested using trypsin, and the resulting peptides were eluted and dried prior to analysis on an Orbitrap Lumos mass spectrometer (Thermo-Fisher). Loading solvent was 3% MeCN, 0.1% FA, analytical solvent A: 0.1% FA and B: MeCN + 0.1% FA. All separations were carried out at 55°C. Samples were loaded at 5 µl/min for 5mins in loading solvent before beginning the analytical gradient. The following gradient was used: 3-40% B over 29mins followed by a 3min wash at 95% B and equilibration at 3% B for 10mins. The following settings were used: MS1: 300-1500 Th, 120,000 resolution, 4 × 10^5^ AGC target, 50 ms maximum injection time. MS2: Quadrupole isolation at an isolation width of m/z 1.6, HCD fragmentation (NCE 35) with fragment ions scanning in the Orbitrap from m/z 110, 5 × 10^4^ AGC target, 60ms maximum injection time, ions accumulated for all parallelisable time. Dynamic exclusion was set to +/- 10 ppm for 60s. MS2 fragmentation was trigged on precursors 5 × 10^4^ counts and above. Data was analysed in MaxQuant version 2.0.1.0, and the method of Significance A was used to estimate *P* values (Cox and Mann, 2008). Full dataset is provided in supplemental files.

### Protein preparation and western blotting

IFITM3 western blots using human cells were performed as described in (Makvandi-Nejad et al., 2018; Wellington et al., 2021). For other proteins, 1 × 10^6^ cells per condition/cell line were homogenized using RIPA lysis buffer (Thermo-Fisher). Protein extracts were prepared for gel electrophoresis by addition of 1:1 Novex^TM^ Tris-Glycine SDS Sample Buffer (Thermo-Fisher) and heating at 85°C for 5min, followed by addition of sample to 12% or 8-16% (dependent on predicted band size) Novex wedgewell Tris Glycine mini gels (Thermo-Fisher) for electrophoresis. Proteins were blotted onto PDVF membrane using the mini-gel wet-transfer XCell II Blot Module (Thermo-Fisher) in transfer buffer (20% methanol, 25mM Tris-base and 192mM Glycine). Membranes were blocked with 5% milk powder in TBS-T (Sigma-Aldrich), then transferred to primary antibody for anti-Nogo-B (0.2μg/ml; Bio-Techne), anti-GAPDH (1μg/ml; Merck Millipore), anti-IFITM1 (3μg/ml; Proteintech), anti-IFITM2 (3μg/ml; Proteintech) or anti-Nogo-A (1μg/ml; Abcam) in 5% milk/TBS-T and incubated overnight at 4°C. Primary antibodies were probed with IRDye 680LT goat anti-mouse (Li-Cor, 926-68020), IRDye 800LT goat anti-rabbit (Li-Cor, 925-32210) or Alexa Fluor 680 donkey anti-sheep IgG (H+L; Invitrogen), and visualized using the Li-Cor Odyssey Imaging System. Protein band intensity was determined using ImageJ, with values calculated relative to GAPDH with background removed for the same sample lane. In murine cells, 1 × 10^5^ cells per condition/cell line were homogenized using 1 × NuPAGE^TM^ LDS sample buffer (Thermo-Fisher) supplemented with 100mM dithiothreitol (DTT) (Sigma-Aldrich) and boiled at 100°C for 10min. Samples were run using a NuPAGE^TM^ 4 to 12% Bis-Tris gels (Thermo-Fisher) for electrophoresis. Proteins were blotted onto PDVF membrane using the mini-gel wet-transfer XCell II Blot Module (Thermo-Fisher) in transfer buffer (1 × NuPAGE^TM^ Transfer Buffer (Thermo-Fisher)). Membranes were blocked with 5% milk powder in PBS-T (Sigma-Aldrich), then transferred to primary antibody for anti-Nogo-B (0.2μg/ml; Bio-Techne), anti-fragilis (IFITM3, 1μg/ml; Abcam) anti-Ubiquitin (1μg/ml; Abcam), and anti-ACTIN (1μg/ml; Sigma-Aldrich) in 0.5% milk/PBS-T and incubated overnight at 4°C. Primary antibodies were probed with either rabbit-anti-sheep IgG (H+L)-HRP conjugate, goat anti-rabbit IgG (H+L)-HRP conjugate (both Biorad) or Veriblot IP detection reagent (Abcam) and visualised using a Syngene G:Box imaging system. Protein band intensity was determined using ImageJ, with values calculated relative to ACTIN with background removed for the same sample lane.

### Immunostaining and imaging

iPS-DCs were harvested from plates, washed in PBS, fixed in 4% PFA for 10min at room temperature, and then spun onto 0.01% poly-L-lysine (Sigma-Aldrich) coated slides using a Cytospin cytocentrifuge. Immediately after spinning onto slides, residual aldehydes were quenched using 25mM glycine for 10min, followed by three washes with PBS. Cells were permeabilised with 0.5% saponin (Fisher Scientific), followed by blocking with 0.05% saponin and 2% BSA (Sigma-Aldrich) for 1h, and incubation with primary antibodies for anti-TLR2 (1:100; Abcam) and anti-Nogo-B (BioTechne) diluted 1:100 in blocking solution for 1h Samples were washed with 0.05% saponin three times, and then fluorescently conjugated secondary antibodies (Donkey anti-sheep IgG Cy5, Merck Millipore; Rabbit anti-goat IgG FITC, Abcam) diluted in blocking solution were added for 30min. Samples were washed three times with 0.05% saponin and then nuclei were counterstained with DAPI (NucBlue; Invitrogen). Samples were washed three times and then mounted with Vectorshield (Vector Laboratories) and left to dry overnight before imaging on a DeltaVision Elite system, with images captured using a CoolSNAP HQ2 camera. For colocalization analysis, z-stacked images were deconvolved with Huygens deconvolution software (SVI), followed by quantification of colocalization and statistical summarisation using Manders overlap coefficient on ImageJ Fiji.

### THP-1 culture and siRNA

The human monocytic cell line THP-1 was used for siRNA knockdown experiments. THP-1 cells were cultured in RPMI 1640 (Gibco, Thermo-Fisher) supplemented with 10% FBS, 100U/mL Penicillin/streptomycin and 2mM L-glutamine (Gibco, Thermo-Fisher). For siRNA knockdown, THP-1s were transferred to Accell siRNA Delivery medium (Horizon Discovery). Cells were targeted with 1μM non-targeting siRNA and/or SMARTPool directed against *RTN4* or *IFITM3* (Horizon Discovery). 72h post-targeting, THP-1s were plated for assays at 1 × 10^5^ cells per condition per replicate. Knockdown was assessed at 72h post-targeting by western blot for IFITM3 and Nogo-B expression, as described above. For murine BM-DC siRNA transfection, cells were differentiated as described above. During differentiation (d7), 1 × 10^6^ BM-DCs were transfected with 150nM siRNA targeting either *Rtn4* or control siRNA (Mus musculus flexitube siRNA (*Rtn4*) or AllStars negative control siRNA (control)) (Qiagen), using HiPerfect transfection reagent (Qiagen). Cells were left for 24h and then a repeat transfection with siRNAs was performed as before, (d8 of differentiation). Cells were used at d9 of differentiation using 1 × 10^5^ cells per condition per replicate. Knockdown was assessed at 48h post initial transfection by western blot for Nogo-B expression, as described above.

### Flow cytometry

Flow cytometry was performed as described in (Forbester et al., 2020). Briefly, for analysis of surface markers on iPS-DCs, cells were stained with Zombie Aqua fixable dye (BioLegend), Fc receptors were blocked using human TruStain FcX (BioLegend), and cells were then subsequently stained for surface markers anti-CD11c-FITC (Bu15, Biolegend) and anti-CD141-APC (M80, Biolegend), followed by fixation with 4% paraformaldehyde. For the detection of IAV nucleoprotein, cells were stained with Zombie Aqua fixable dye, fixed with 4% paraformaldehyde, and permeabilized with 0.5% Triton X-100, followed by incubation with human TruStain FcX and staining with anti-influenza A virus nucleoprotein antibody (431; Abcam) in 0.1% Triton X-100 solution. Data were acquired using an Attune NxT flow cytometer (Thermo-Fisher). Electronic compensation was performed with antibody (Ab) capture beads (BD Biosciences) stained separately with individual monoclonal antibodies (MAbs) used in the experimental panel. Data were analyzed using the FlowJo software (TreeStar, Inc.).

### Statistical analyses

Statistical significance was performed using the GraphPad Prism software. The Mann-Whitney U or Student’s *t* test was used for two-group comparisons. To analyse data from multiple groups, 2-way ANOVA analysis was performed. Mass Spectrometry statistical *P* values were estimated using Benjamini-Hochberg-corrected significance A values (Cox and Mann, 2008). A *P* value of ≤0.05 was considered to be significant. For all tests performed, *P* values are reported as follows: n.s., >0.05; *, ≤0.05; **, ≤0.01; ***, ≤0.001; and ****, ≤0.0001.

## Acknowledgments

This work was funded by a Wellcome Trust Senior Research Fellowship to I.R.H. (grant 207503/Z/17/Z); a Wellcome Trust Senior Research Fellowship to M.P.W. (grant 108070/Z/15/Z); the Medical Research Council, United Kingdom (grant MR/L018942/1 and MRC Human Immunology Unit Core); and the Chinese Academy of Medical Sciences (CAMS) Innovation Fund for Medical Sciences (CIFMS), China (grant 2018-I2M-2-002). Pragati Sabberwal was funded by a Cardiff University Systems Immunity University Research Institute PhD Studentship. The Wellcome Trust Sanger Institute was the source of the Kolf2 human induced pluripotent cell line, which was generated under the Human Induced Pluripotent Stem Cell Initiative funded by a grant from the Wellcome Trust and the Medical Research Council, supported by the Wellcome Trust (grant WT098051) and the NIHR/Wellcome Trust Clinical Research Facility. IFITM3 iPSC knockout lines were generated, characterised and banked by the Gene editing core facility at the Wellcome Sanger Institute. The graphical abstract was drawn using bioRENDER.

## Ethics statement

All animal studies were performed at Cardiff University (Heath Park research support facility) under UK Home Office Project License number P7867DADD, as approved by the UK Home Office, London, United Kingdom. Written consent was obtained for the use of cell lines for the HipSci project from healthy volunteers. Ethical approval was granted by the National Research Ethics Service (NRES) Research Ethics Committee Yorkshire and The Humber-Leeds West, under reference number 15/YH/0391.

## Author contributions

M.C., J.L.F., M.M., P.S., D.W., S.D., S.C., K.H., L.N., R.A., B.J., M.C., S.M-N., and J.A.L., performed experiments; M.C., J.L.F., D.W., L.N., R.A. M.P.W., R.J.S., T.D., and I.R.H., analysed the data; S.M.S., T.D., M.P.W., and R.J.S., provided reagents; I.R.H directed the study; M.C., J.L.F., S.M.S., and I.R.H wrote the manuscript.

---

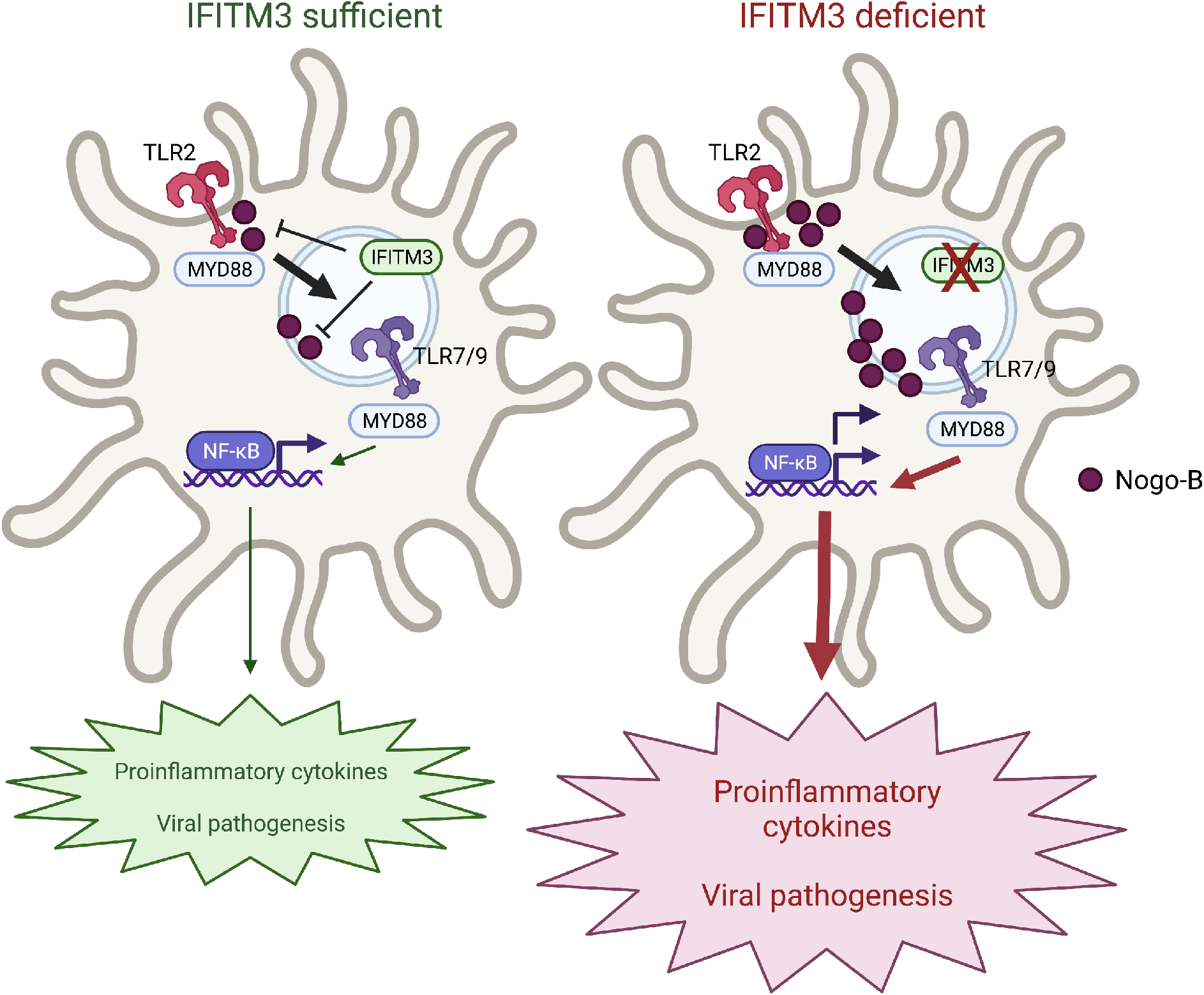

