## Supplemental figures for "IFITM3 regulates virus-induced inflammatory cytokine production by titrating Nogo-B orchestration of TLR responses"

### Supplemental Figure 1

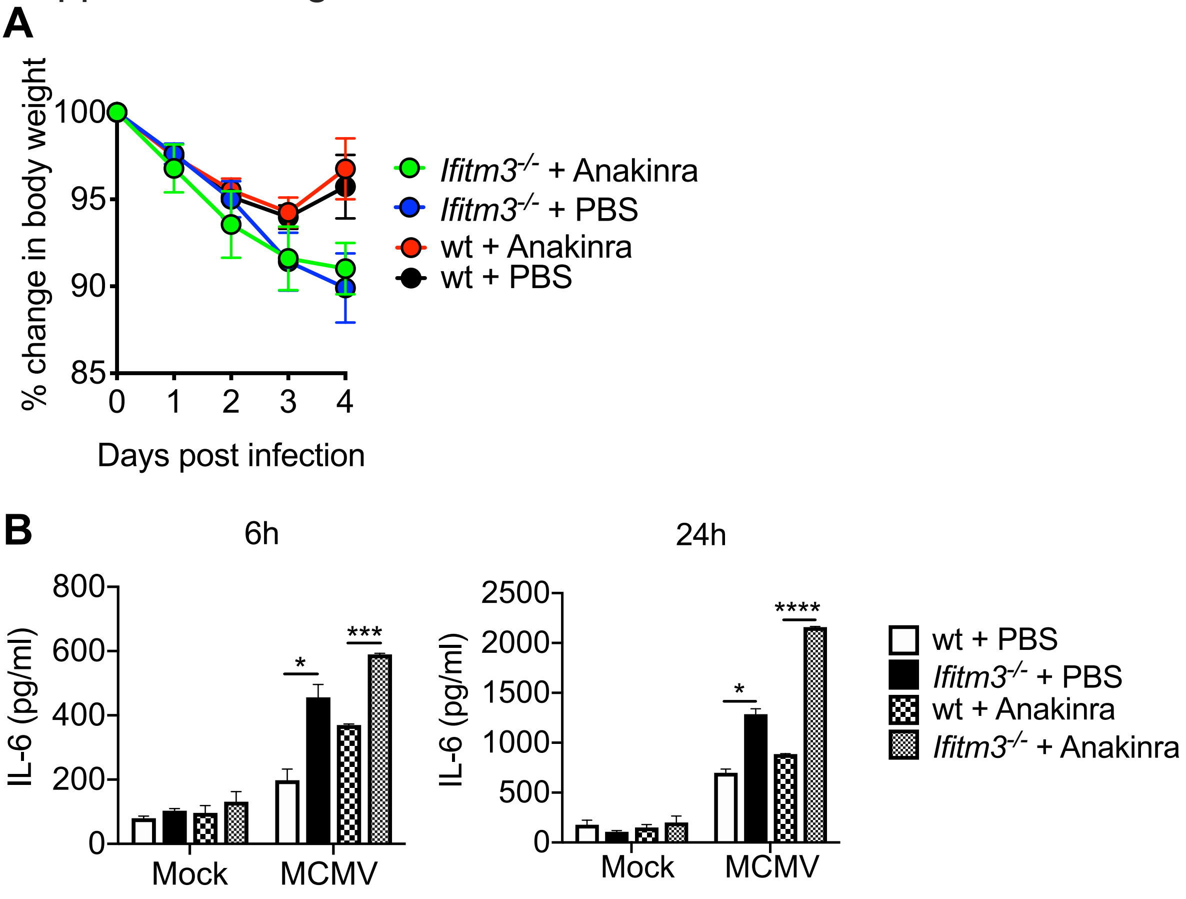

Supplemental Figure 2

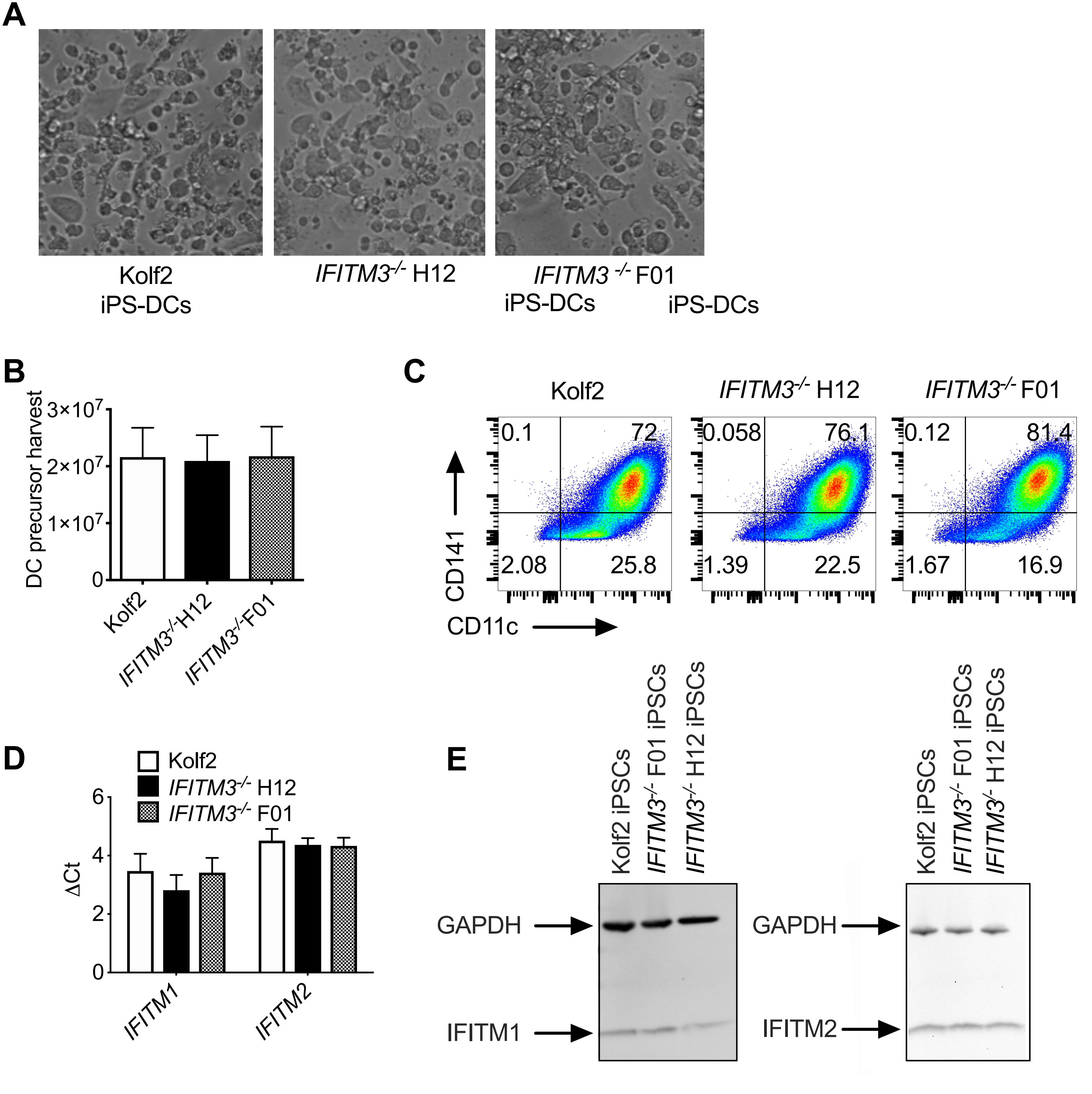

### Supplemental Figure 3

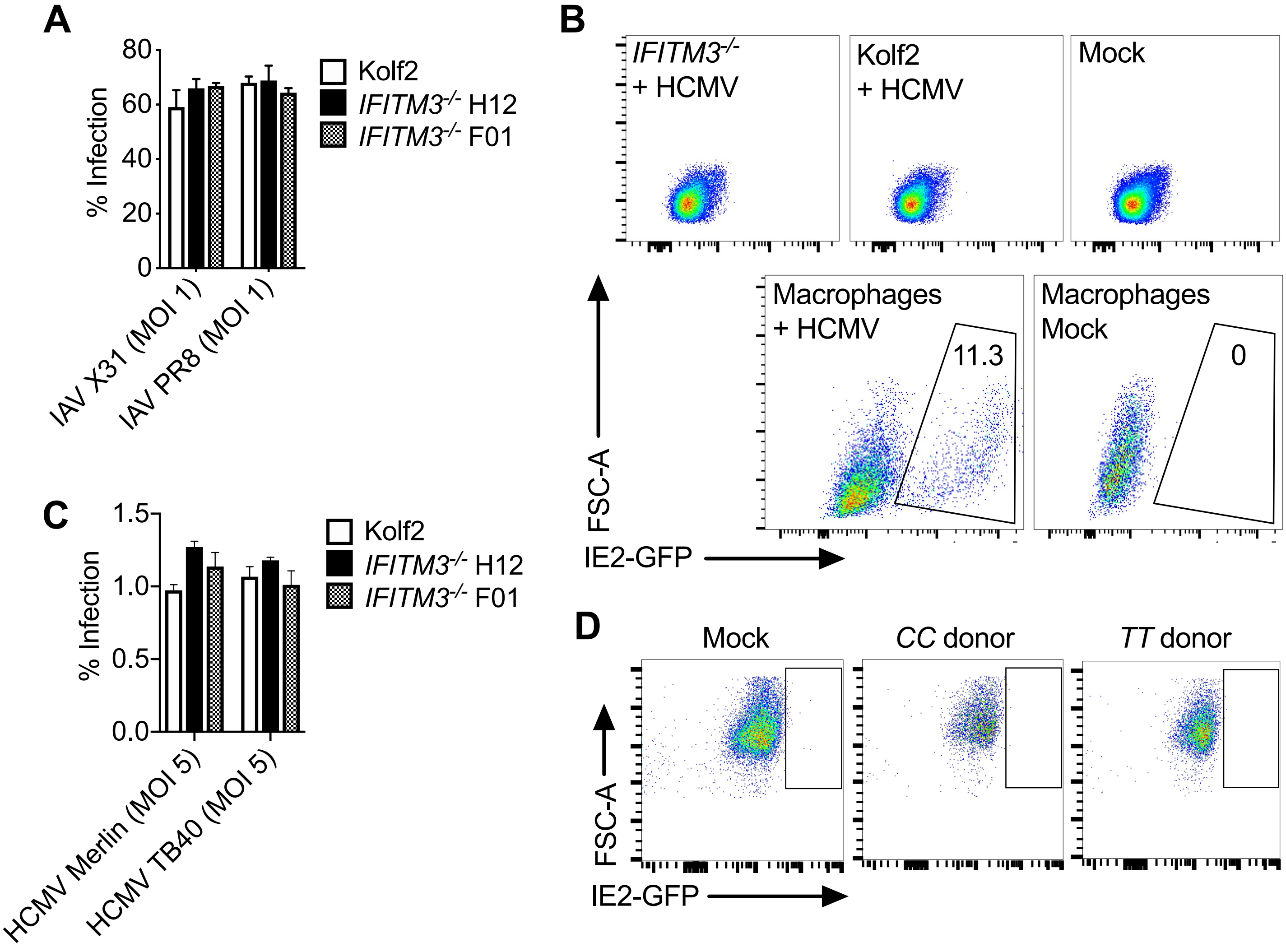

### Supplemental Figure 4

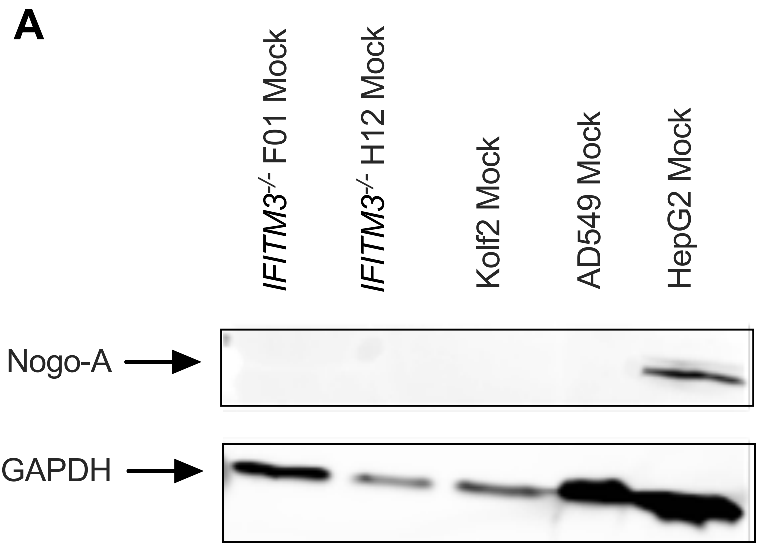

### Supplemental Figure 5

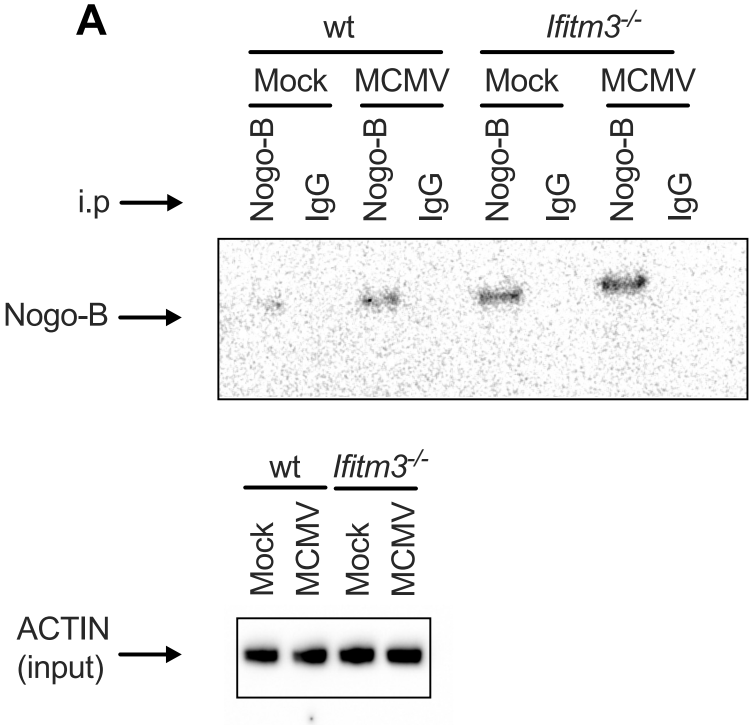

### Supplemental Figure 6

**A**

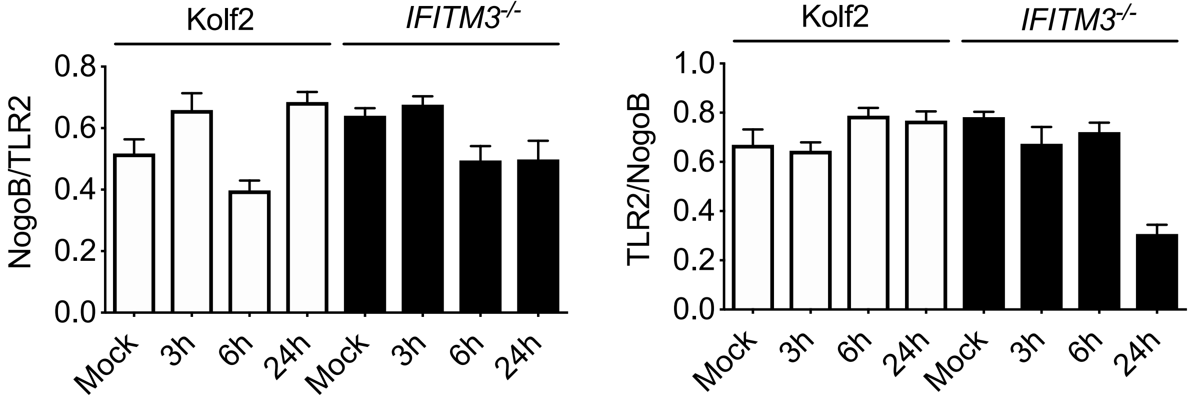
